# Characterizing homology-induced data leakage and memorization in genome-trained sequence models

**DOI:** 10.1101/2025.01.22.634321

**Authors:** Abdul Muntakim Rafi, Brett Kiyota, Nozomu Yachie, Carl de Boer

## Abstract

Models that predict function from DNA sequence have become critical tools in deciphering the roles of genomic sequences and genetic variation within them. However, traditional approaches for dividing the genomic sequences into training data, used to create the model, and test data, used to determine the model’s performance on unseen data, fail to account for the widespread homology within genomes. Using simulations, we illustrate how homology-based data leakage can lead to overestimation of model performance. Across a variety of genomics models, we demonstrate that performance on test sequences varies systematically by their similarity with training sequences. Models generally perform well on distant sequences, reflecting the application of learned generalizable principles. At higher and intermediate similarity, models rely on memorized associations, inflating performance when function is conserved between homologs but failing when homologous sequences have functionally diverged. To dissect and mitigate these effects, we introduce hashFrag, a scalable solution for homology detection and data partitioning. Using hashFrag, we demonstrate how to create homology-aware evaluations of model performance, and improve model generalizability by providing improved splits for model training. Altogether, we establish how homology creates a systematic bias in genome-trained models and must be accounted for to ensure reliable evaluation of sequence-to-function predictors.

## Introduction

Machine learning has emerged as a powerful tool for understanding how the sequence of the genome encodes its function (1–4). Many sequence-to-function models aim to learn cis-regulatory logic, namely how transcription factor binding sites, their combinatorial syntax, and chromatin context work together to control gene expression and cellular identity (5–12). These models have already enabled the design of cell type-specific regulatory sequences (13–17) and are increasingly used for variant effect prediction (18–22), genome-wide association study (GWAS) fine-mapping (23), and clinical interpretation of noncoding mutations (24–26).

Learning causal, context-dependent cis-regulatory logic from genomic data is, however, fundamentally constrained by the nature of the genome itself (27–29). Many models are trained on genome-wide chromatin profiling assays (11,18,30–36) or on fixed-context reporter assay measurements of genomic sequences (13,37–40). However, nearly half the human genome consists of repeat elements (41), and many sequences have homologs across different chromosomes often in multiple copies, resulting from transposition, retroposition, translocation, whole genome duplication, and other processes that permeate genomes (42). How this repetitiveness impacts the behavior of models trained on the genome remains poorly characterized (43,44).

In machine learning, data are typically divided into a training set used to tune model weights, a validation set used to determine when the model is most generalizable and training should be stopped, and a test set, held out from training, that is used to estimate how well the model will generalize to new examples (45). Data leakage, defined as the presence of non-independent information across data splits, can bias model evaluation. Leakage between train and validation sets causes underestimation of overfitting, resulting in suboptimal model selection, while leakage between training and test sets inflates performance estimates (46). In genomics, such leakage can arise when homologous sequences span data splits, causing related sequences to appear in multiple sets (27).

In this study, we systematically quantify how homology distorts both training and evaluation of sequence-to-function models. Using controlled simulations, we first show that homology between training and test sequences can result in suboptimal model selection and performance overestimation. We then analyze how homology impacts performance for three representative sequence-to-function modeling scenarios: fixed-context MPRA predictors (200 bp) (39,47), short-context ChromBPNet (2 kb) (33), and long-context Enformer (196 kb) (31). Across all models, we find that predictive accuracy across test sequences varies systematically with similarity to training sequences. While models generalize well to distant sequences, performance at higher similarity is inconsistent: memorization of training examples inflates accuracy when function is conserved but degrades it as function diverges. This effect is particularly severe in long-context models like Enformer (31), where prediction accuracy drops to its lowest point at high similarity as locally similar sequences in different contexts differ in their functions. These observations are consistent with genomics models fitting the training data by simultaneously learning general rules of genome regulation and memorizing trends that often fail to generalize. Because homology is pervasive and inherent to genome structure, this bias is not specific to any particular model or architecture but is a fundamental property of genomic data that affects all genome-trained sequence-to-function models. To address this challenge, we introduce hashFrag, a scalable solution leveraging Basic Local Alignment Search Tool (BLAST) (48,49) to detect sequence homology. We demonstrate how hashFrag can be used to stratify test sets to quantify homology-dependent performance, filter homologous sequences for reliable generalization estimates, and create homology-aware splits that improve model training.

## Results

### Simulation reveals inflated performance with homology-based data leakage

We performed a controlled simulation (**Fig. 1A**) to investigate the effect of homology on model evaluation. We created positive and negative sequence sets by randomly generating 200 bp sequences and inserting 1–3 out of a possible 20 transcription factor binding sites (TFBSs; each 10-bp long) randomly into positive set sequences, while ensuring negative sequences had no TFBSs. We then generated 20 homologous variants for each starting sequence by introducing random point mutations at a 5% rate, restricted to avoid the inserted TFBSs. We then split the data into separate sets for training and testing. The pooled sequences were split into training and test sets, and the test set was stratified into two subgroups: "leaked" sequences that share homologs with the training set, and "non-leaked" sequences that do not. We then trained convolutional neural networks (CNNs) of increasing capacity (Shallow, Medium, High-Capacity; **Methods**) to predict the classes (presence or absence of TFBSs) from sequence alone (**Fig. 1A; Methods**).

**Figure 1:**
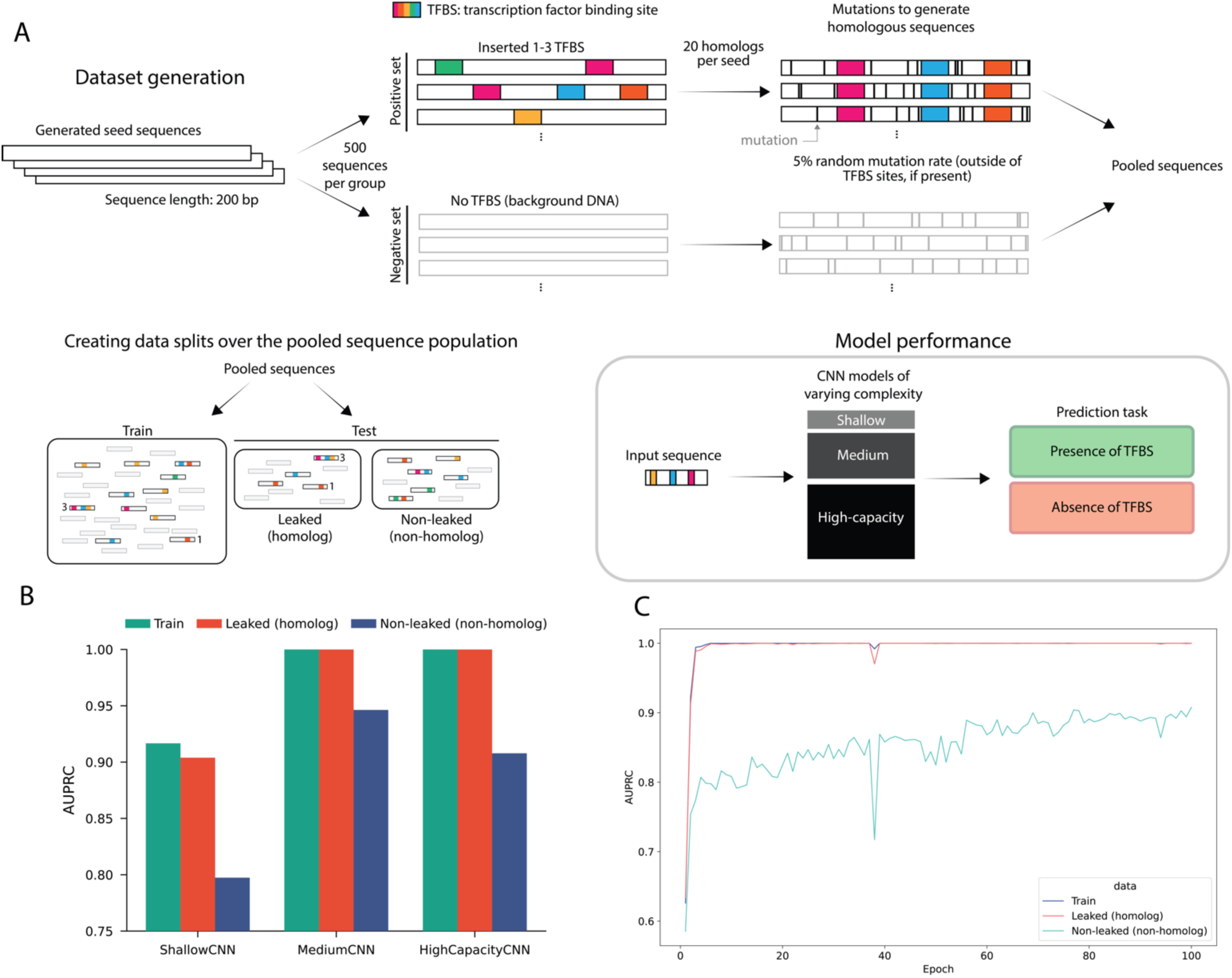
Homology-based data leakage systematically inflates model performance estimates in controlled simulations. **(A)** Schematic of synthetic dataset generation and training models to predict presence or absence of TFBSs. **(B)** Model performance (AUPRC, *y*-axis) on train, leaked test, and non-leaked test subgroups (colors) across three model architectures (*x*-axis). **(C)** Performance (AUPRC, *y*-axis) tracked during training (epochs, *x*-axis) for HighCapacityCNN model on training, leaked test, and non-leaked test subgroups (colors).

Across all architectures, leaked test sequences had systematically higher apparent performance than non-leaked test sequences (**Fig. 1B**). Moreover, at the highest model capacity, the performance gap between leaked and non-leaked test subsets widened, consistent with higher-capacity models being more capable of memorizing, which can hurt model generalization (50,51). We also monitored model performance during training (**Fig. 1C**). The model quickly converged for leaked test sequences, obtaining near-perfect performance on both training and leaked sequences within 5 epochs of model training. In contrast, performance on non-leaked test sequences converged slowly with longer training (**Fig. 1C**). Together, this demonstrates that homology-based leakage alone is sufficient to create inflated performance metrics (46,52) and a leaky validation set, can lead to poor model selection by failing to select the optimal model (46,53).

### Potential impact of homology on sequence-to-function models and variant effect prediction

Homologous sequences distributed across chromosomes can share the same signal. For example, two homologous 1,000 bp regions on chromosomes 9 and 16 in K562 cells exhibit nearly identical ATAC-seq read count patterns (**Fig. 2A**), despite their distinct genomic locations. When such homologous regions are split across training and test sets, models can achieve high performance by memorizing the signal without learning the underlying regulatory logic.

**Figure 2:**
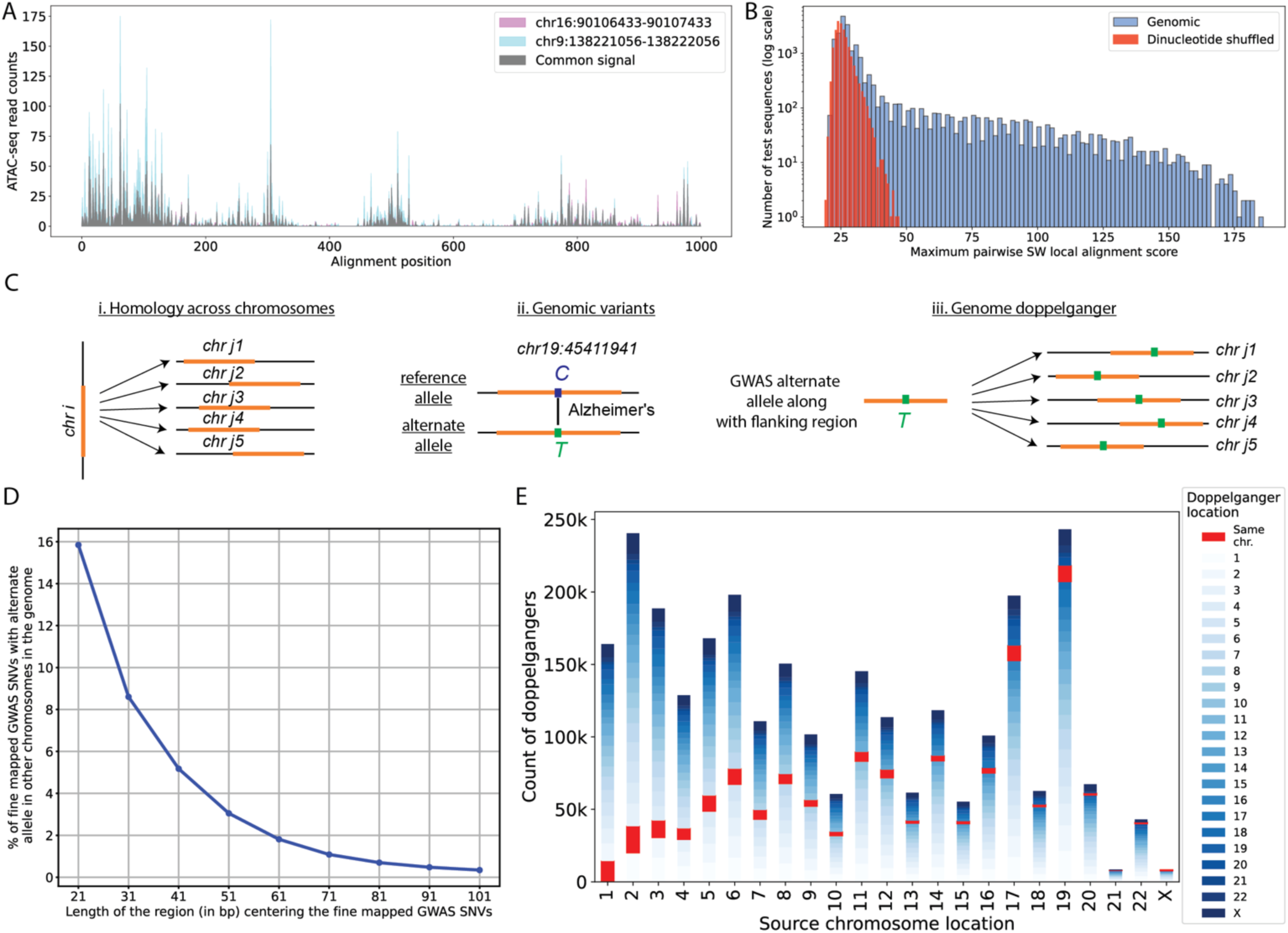
Potential for homology-based data leakage in sequence-to-function models trained on genomic DNA sequences. (**A**) Homologous sequences from two different chromosomes can share functional genomic signals. ATAC-seq read counts (*y*-axis) for two homologous 1000 bp regions (*x*-axis) on chromosomes 9 and 16 (colours) in K562 cells. (**B**) Homology is common between chromosomes. Histogram showing the number of test sequences (*y*-axis) with corresponding maximum pairwise SW local alignment scores with the training sequences (*x*-axis) for both genomic (blue) and dinucleotide shuffled (red) sequences, with training and test sets randomly sampled from distinct chromosome sets (20,000 sequences each). (**C**) Illustration of (i) homology across chromosomes, (ii) SNVs associated with diseases, and (iii) SNV doppelgängers, sequences elsewhere in the genome with an identical sequence to the GWAS alternate allele, including its flanking region. (**D**) Percentage of GWAS SNVs (*y*-axis) with SNV doppelgängers of each sequence length (*x*-axis). (**E**) The number of 41 bp SNV doppelgängers (*y*-axis) that are located on each chromosome (colours), for fine-mapped SNVs originating from each chromosome (*x*-axis).

To quantify the extent of cross-chromosomal homology, we analyzed putative enhancers from K562 cells that were used in an MPRA experiment (39). We divided the sequences into two halves by chromosome and calculated the maximum pairwise Smith-Waterman (SW) local alignment scores (54) for sequences between these two sets. SW local alignment is well suited for detecting shared ancestry because it rewards contiguous tracts of sequence identity and conservative substitutions that arise through evolutionary descent (**Supplementary Figure 1**). Importantly, convergently evolved regulatory sequences, such as independently acquired transcription factor binding sites that share a motif but not ancestry (55), are expected to receive low SW scores, since short, dispersed motif matches do not produce the extended conservation required to yield high local-alignment scores. Many genomic sequences exhibited substantially higher similarity than expected from dinucleotide-shuffled control sequences (**Fig. 2B, Supplementary Figure 2**), indicating that sequence similarity between chromosomally separated training and test sets extends well beyond what would arise by chance. Many sequences fell into distinct clusters of homology, with many members in each chromosome split (**Supplementary Figure 3**). Consistent with about half of the genome being derived from repeat elements (41), we found that nearly half of these putative K562 regulatory elements overlap with annotated repeat elements (**Supplementary Figure 4**). to have higher alignment scores to sequences on other chromosomes compared to non-repeat sequences (**Supplementary Figure 4**). This suggests that a substantial fraction of putative regulatory elements used in model training may be subject to homology-based leakage under standard chromosomal splitting approaches.

Homology can also impact the interpretation of genetic variation, one of the critical applications of sequence-to-function models. To enrich for variants with phenotypic consequences, we restricted our analysis to single nucleotide variants (SNVs) identified by genome-wide association study (GWAS) variants with posterior inclusion probabilities (PIPs) greater than 0.1 (56). A substantial percentage of these high-confidence GWAS SNVs have their alternate alleles, along with flanking sequences, replicated on other chromosomes, effectively creating "SNV doppelgängers" (**Fig. 2C, 2D**). Many SNV doppelgängers appear multiple times throughout the genome (**Fig. 2E**, **Supplementary Figure 5**). Doppelgängers of 41 bp (20 bp flanking each side of the SNV; **Fig. 2E**) are sufficiently unique (∼1 in 5×10²⁴ possible sequences) that an exact match elsewhere in the genome almost certainly reflects true homology rather than chance, and are long enough for a model to memorize a specific activity associated with that sequence. This creates a fundamental challenge for model evaluation. Since many alternate allele sequences appear repeatedly in the training data, predictions may stem from memorized sequence-function relationships rather than learned cis-regulatory logic (57,58).

### Empirical evidence of homology-based leakage in MPRA models

While one could exclude repeat-derived regulatory elements during model training, this would halve the available data and incompletely eliminate homology-based leakage. In practice, repeat elements are downweighted when training DNA language models to prevent learning trivial copy-paste patterns (59–62). However, applying this strategy would be problematic for sequence-to-expression modeling because many functional elements are derived from repeats (63–65). Rather than excluding homologous sequences, we sought to ascertain the degree to which overfitting to homologous training sequences could be affecting model performance.

We asked how good a performance could be achieved by a model that relies exclusively on memorization of homologous training sequences, without any understanding of cis-regulatory logic. As a maximally overfit benchmark, we developed a simple heuristic model, OverfitNN, which predicts gene expression by averaging the expression labels of the nearest neighbor sequences from the training set (**Fig. 3A, Methods**). We evaluated OverfitNN across test sequences, stratified into bins by their similarity to training sequences (maximum SW alignment score). OverfitNN was able to predict expression reasonably well for highly similar sequences, but its performance steadily dropped as the test sequences became less similar to its training set (**Fig. 3B, Supplementary Figure 6**). Next, we compared OverfitNN to several state-of-the-art neural networks, including different model architecture types (DREAM-CNN (66), DREAM-RNN (66), DREAM-Attn (66), and MPRAnn (39)), trained them on these MPRA data (39) using the same chromosomal train-test splits (**Methods**). If a model has learned all regulatory mechanisms equally well, it there should be no relationship between performance on test sequences and their similarity to training sequences. If, however, regulatory mechanisms were learned to different degrees, the performance-similarity relationship is expected to be monotonic and positive, as performance is better as sequences are more similar to training data (and therefore share more regulatory mechanisms). In contrast, every model tested varied in their performance on test sequences by their similarity to training sequences, but with a U-shape rather than monotonic or flat **(Fig. 3B, Supplementary Figure 6)**. Interestingly, we found that the performance on the test sequences that are most similar to the training data is actually superior to the performance on the training data itself early in model training (**Fig. 3C, Supplementary Figure 7**), consistent with homology-based overfitting such as that we observed in our controlled simulation (**Fig. 1C**). The inferior model performance at intermediate similarity levels (e.g. performances at intermediate scores of 50-100 are consistently worse than performance at low scores of 30-50; **Fig. 3B, Supplementary Figure 6**) likely reflects the models’ dependence on homology backfiring as the regulatory sequences diverge to the point where homology to training sequences can be detected but is not helpful in making predictions. Together, these trends suggest that models rely on memorized associations that are rewarded at very high similarity, where sequence conservation entails functional conservation. At intermediate similarity, however, the same memorization strategy becomes unreliable as sequences retain enough homology to trigger memorization-based predictions yet have diverged sufficiently that sequence similarity no longer implies functional similarity, actively misleading the model. Performance recovers at the lowest similarity levels, where the absence of detectable homology forces models to use learned regulatory features rather than memorized associations.

**Figure 3:**
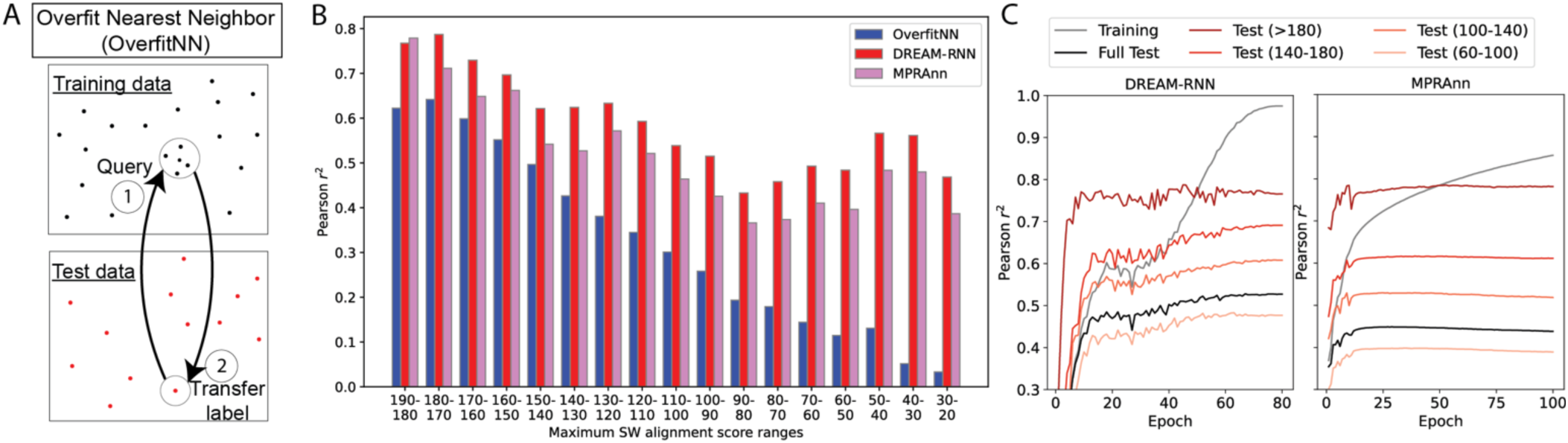
Evidence of homology-based data leakage in neural networks trained on MPRA data with genomic DNA sequence library. (**A**) Schematic of OverfitNN, which predicts gene expression using sequence similarity rather than cis-regulatory logic. For each test sequence, the nearest neighbor in the training set is identified by SW alignment (step 1), and its expression label is transferred as the prediction (step 2). (**B**) Performance comparison (Pearson 𝑟^2^; *y*-axis) of models (OverfitNN, MPRAnn, and DREAM-RNN; colors) across varying levels of sequence similarity (maximum SW alignment score, *x*-axis). (**C**) Neural networks trained on chromosomal splitting (chromosomal split 1, Methods) exhibit varying performance levels (Pearson 𝑟^2^; *y*-axes) on unseen data during model training (*x*-axes) depending on the degree of homology-based leakage (colors).

### hashFrag: a scalable solution for detecting and mitigating homology-based leakage

Detecting homology systematically presents a substantial computational challenge due to the quadratic scaling of pairwise SW alignment score computation time with respect to both sequence length and dataset size. Consequently, exhaustive pairwise comparison is infeasible for longer sequences and larger datasets typically used by many genomic models (30,18,31,34,10,12). To create a scalable solution for characterizing homology within genomics training data, we developed hashFrag, a computational tool that circumvents the computational complexity of calculating pairwise local alignments by leveraging the heuristics and optimizations in BLAST. hashFrag initially constructs a BLAST database of all sequences in the dataset, and then queries each sequence against the database to shortlist candidate homologous sequences, which can subsequently be assessed for homology based on their alignment scores (**Fig. 4A**). We applied hashFrag to the MPRA dataset (39) and demonstrated that it detects homology with near-perfect recall (99.99%) and a false-positive rate (FPR) of ∼0.17% at a homolog-defining SW local alignment score threshold of 60 (**Fig. 4B**). We have two modes in hashFrag: hashFrag-lightning, which uses the heuristic alignment scores provided by BLAST, and hashFrag-pure, which calculates exact SW alignment scores for the candidate pairs. Both versions are substantially faster than naive alternative of computing all pairwise SW alignments exhaustively, with 161-fold speedup for lightning and 86-fold for pure (**Fig. 4C**), enabling partitioning even large sequence sets in the span of a day on a single CPU.

**Figure 4:**
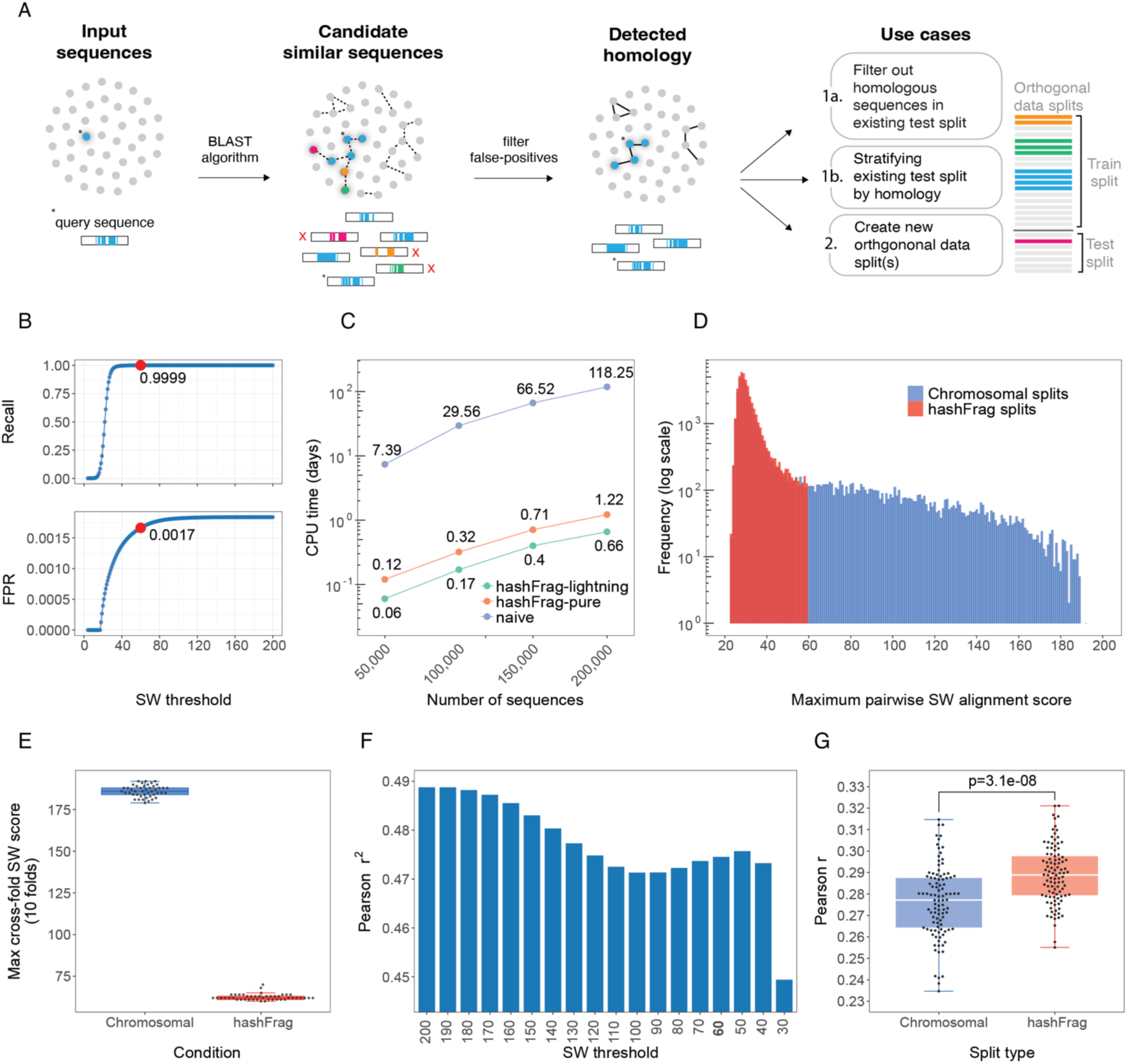
hashFrag as a scalable approach to detect homology-based data leakage. (**A**) Overview of hashFrag. Sequences are queried against a BLASTn database to identify candidate homologs, which are filtered by SW alignment score. Detected homology can be used to filter test sets, stratify by similarity, or create orthogonal splits. (**B**) hashFrag’s recall (*y*-axis; top) and false-positive rate (*y*-axis; bottom) at varying SW thresholds (*x*-axes); the selected threshold of 60 is highlighted in red. (**C**) Runtime (*y*-axis) comparison of naive pairwise SW alignments *vs.* hashFrag (colours) across dataset sizes (*x*-axis). (**D**) Distribution of maximum pairwise SW scores between train and test sets for chromosomal splits (blue) versus hashFrag-pure (red) at an 80/20 split ratio. (**E**) Maximum pairwise SW scores across 10 data folds for chromosomal splits (scores >179) versus hashFrag-lightning splits (scores <70), demonstrating effective orthogonal partitioning. (**F**) DREAM-RNN performance (*y*-axis) after removing test sequences exceeding each SW threshold (*x*-axis). (**G**) DREAM-RNN Performance on designed sequences from Gosai et al. (13) (*y*-axis) when trained on chromosomal or hashFrag train/validation splits (*x*-axis) across 100 replicates (points). P-value: Two-sample, two-sided t-test.

We next validated that hashFrag successfully eliminates train-test leakage where chromosomal splitting fails. While standard chromosomal splits retain high-similarity sequences between training and test sets, hashFrag effectively removes these overlaps (**Fig. 4D**). We also note while hashFrag-pure strictly maintains inter-fold similarity below the specified threshold, hashFrag-lightning occasionally permits a small number of sequences with scores slightly above the threshold (**Supplementary Figure 8**) due to its reliance on heuristic BLAST scores. Since these near-threshold sequences represent distant homologs, hashFrag-lightning remains a suitable alternative when computational speed is prioritized. hashFrag can also partition sequences into multiple leakage free subsets, with minimal similarity for even the most similar sequences in hashFrag folds, whereas all chromosomal folds share very similar sequences (maximum SW alignment score is under 70 for hashFrag and above 175 for chromosomal splits; **Fig. 4E**).

By identifying homology-based leakage, hashFrag can improve our understanding of model generalization and enable better models to be created. Stratifying existing test sets can reveal how performance varies across different similarity ranges (**Fig 3B**), or homologous sequences can be removed from existing test sets, enabling a better estimation of model generalization (**Fig. 4F**, **Supplementary Figure 9**). We hypothesized that partitioning data with hashFrag could result in better model performance because leakage between validation and training data will cause one to underestimate the degree of overfitting during training (**Fig. 1C**), leading to poor generalizability (46,53,52). However, comparing performance of hashFrag and chromosomal-split models on genomic sequences is inherently unfair because the models trained on chromosomal splits will have been told the answer for some test sequences, having seen similar sequences during training. Accordingly, we compared hashFrag and chromosomal-split models on designed sequences from Gosai et al. (13), confirming that they were not similar to any of the genomic sequences used for training (**Supplementary Figure 10**). hashFrag-trained models outperformed chromosomal split-trained models (**Fig. 4G**, **Supplementary Figure 11, Methods**), showing that chromosomal splitting not only introduces train-test leakage but also creates inferior train-validation splits that translate to inferior models.

### Homology memorization confounds generalization in genomic profile trained models

Notably, our analysis of models trained on MPRA data likely underestimates the extent of homology-based leakage in genome-trained models. MPRAs capture the inherent activity of each DNA sequence because each sequence is tested in an essentially identical context (67,68). In contrast, the activity of a sequence in the endogenous genome is a product of both itself and of flanking regions (69,70), which provides additional and more complex opportunities for leakage. Accordingly, we next asked what impact homology has on models trained on functional genomic profiles measured in the endogenous genome. Because these sequences are substantially longer than MPRA fragments, we used hashFrag’s HPC module to efficiently compute pairwise homology across the dataset (**Methods**). We first analyzed ChromBPNet, a widely used model that predicts chromatin accessibility profiles and read counts for 1,000 bp sequences (33), and an adaptation of DREAM-RNN to the ChromBPNet task (66). We focused our evaluation only on the read count predictions to assess how homology impacts the prediction of overall signal magnitude rather than accessibility profiles because insertions and deletions between sequences and the interactions of proteins on chromatin would make aligning profiles extremely complex.

As expected, OverfitNN was able to predict accessibility reasonably well for highly similar sequences, rivaling or surpassing the models, but its performance steadily dropped as the test sequences became less similar to its training set (**Fig. 5A**). Unexpectedly, OverfitNN also showed non-trivial performance at the lowest similarity range (score 0-100; **Fig. 5A**), which appears to be explained by the nearest neighbors sharing GC content (**Supplementary Figure 12**), and GC content being correlated with regulatory activity (71–74). ChromBPNet and DREAM-RNN exhibited the same U-shaped performance pattern seen in the MPRA models, with high performance at the extremes of sequence similarity and reduced performance at intermediate levels (SW scores 300-700; **Fig. 5A**). Both models performed well on sequences with little similarity to training data, consistent with the application of learned generalizable principles to sequences distant from the training data (**Fig. 5A**). The drop at intermediate similarity is consistent with sequences being similar enough to trigger memorized associations but functionally diverged enough that those associations no longer hold.

**Figure 5:**
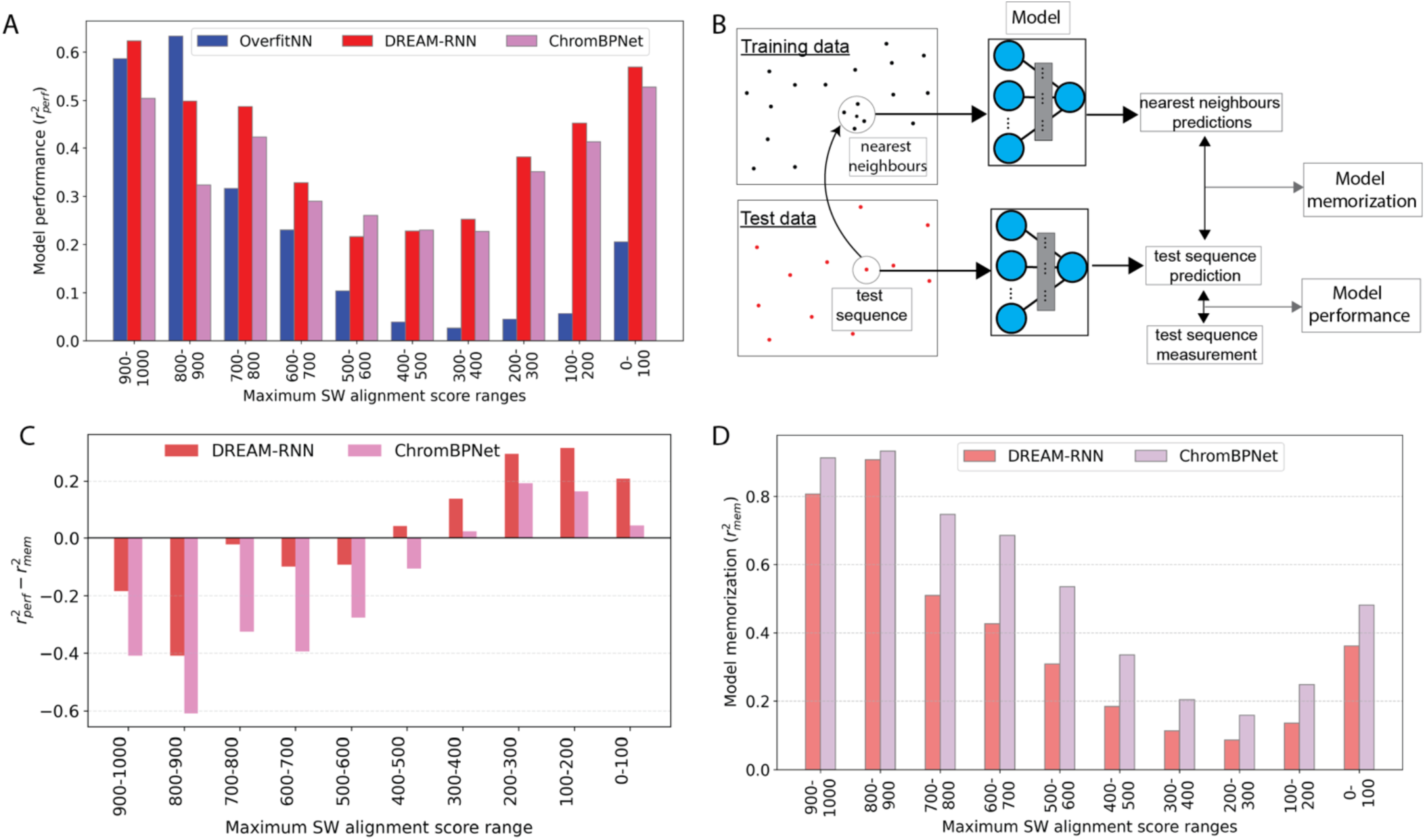
Homology-mediated memorization confounds predictions in genomic profile-trained models. (**A**) Test set read coverage prediction performance (Pearson *r²*; *y*-axis) of OverfitNN, DREAM-RNN, and ChromBPNet (colors) across varying levels of sequence similarity (maximum SW alignment score ranges, *x*-axis) between test and training sequences. (**B**) Schematic of model memorization and performance evaluation. To calculate memorization, for each test sequence, predictions from the 10 most similar training sequences within the maximum SW score range are averaged and compared against the test sequence prediction. To evaluate model performance, the test sequence prediction is compared against the experimental measurement. (**C**) Comparison of gap between model performance and memorization in Pearson *r²* (*y*-axis) for DREAM-RNN and ChromBPNet (colors) across varying levels of sequence similarity between training and test sequences (maximum SW alignment score ranges, *x*-axes). (**D**) Memorization (Pearson *r²*; *y*-axis) of DREAM-RNN and ChromBPNet (colors) across varying levels of sequence similarity between training and test sequences (maximum SW alignment score ranges, *x*-axis).

To investigate the mechanism underlying the performance drop, we next quantified the degree to which the models memorize their training data. Alongside the generalizable rules we want models to learn, models can also learn sequence-specific associations that fail to generalize. Comparing memorization against performance on held-out data lets us distinguish the two. For each test sequence, we identified its nearest neighbors in the training set (by SW score), generated model predictions for those neighbors, and averaged them (**Fig. 5B**). The similarity between this average and the model’s prediction on the test sequence, quantified by Pearson *r²* and mean squared error (MSE), serves as our estimate of memorization. This is analogous to OverfitNN, but uses model predictions in place of experimental measurements for training sequences.

To directly compare memorization and performance, we computed their difference across similarity ranges (**Fig. 5C**). Negative values of *r²_perf_−r²_mem_* indicate that a model’s predictions on test sequences correlate more strongly with its predictions for related training sequences than with the ground truth, suggesting that predictions are better explained by memorization. At higher similarity levels (SW score 500–1000), memorization exceeded performance for both models, consistent with models relying on memorized associations. At intermediate similarity ranges (SW score 500-800), memorization dominated predictions while model performance dropped (**Fig. 5A, 5C**). This suggests that the model continues to rely on memorization-based associations, but because the sequences have diverged, those associations are no longer accurate. Memorization and performance are balanced around SW score 300–500 (**Fig. 5C**), and performance exceeds memorization thereafter as there is increasingly little homology to detect. Further, examining memorization through MSE revealed that model predictions become numerically less anchored to training neighbor predictions as sequences diverge (**Supplementary Figure 13**). The overall gradual transitions from memorization to performance (**Fig. 5C**) is consistent with the models transitioning from memorized to generalizable sequence-activity relationships. Notably, ChromBPNet showed a consistently larger memorization component than DREAM-RNN across all similarity ranges, in both Pearson *r*² (**Fig. 5C,D**) and MSE (**Supplementary Figure 13**), potentially explaining DREAM-RNN’s superior performance (**Fig. 5A**). This illustrates that model architectures and/or training choices can shape the degree to which models memorize, impacting their performance.

### Homology-biased performance in bin-level evaluation of long-range sequence models

We next asked how homology also impacts performance in models that process substantially longer genomic contexts. We analyzed Enformer (31), a transformer-based model whose prediction bins are large enough (128 bp) to identify leakage with hashFrag. Enformer takes 196,608 bp as input and predicts 5,313 human regulatory profiles (DNase-seq, ChIP-seq, CAGE) across the 114,688 bp center region, outputting predictions in 896 bins of 128 bp each (**Fig. 6A**). While leakage would likely be obscured by using a single alignment score for entire 196,608 bp input sequences, we hypothesized that local memorization could be detected if regulatory patterns (e.g. transcription factor binding, open chromatin) (75) are shared between training and test *bins*, regardless of their surrounding context. To rigorously quantify this, we computed the maximum local SW alignment score for all 128 bp test bins against all bins present in the training set, grouped the bins into homology score ranges as before, and calculated model performance and memorization within each group (**Methods**).

**Figure 6:**
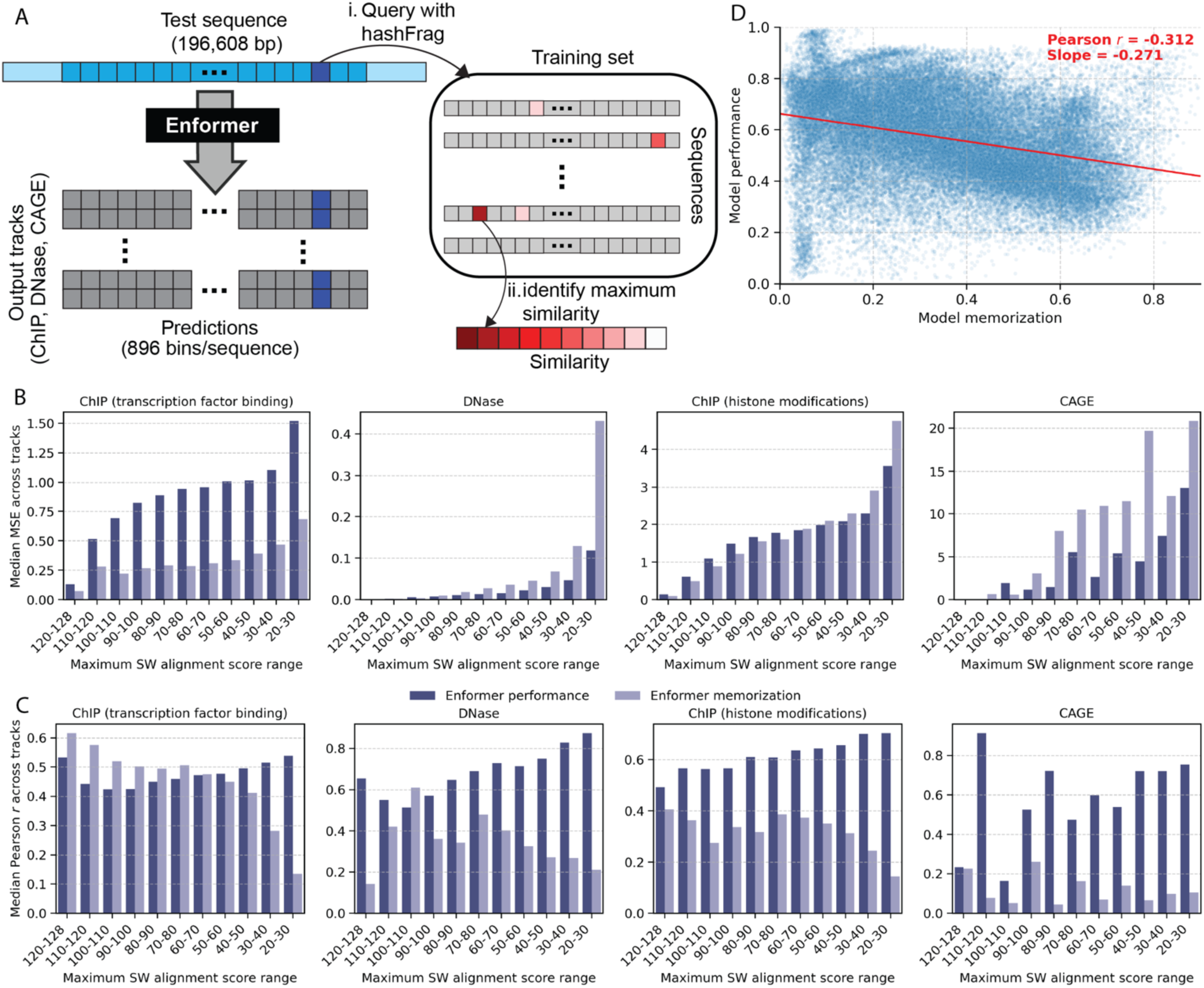
Homology analysis at bin level reveals performance bias in the Enformer model. (**A**) Schematic of the Enformer model’s workflow. The model takes a 196 kb input sequence and predicts expression values for 5,313 human genomic tracks (CAGE, ChIP, DNase) across a center 114 kb region, divided into 896 bins of 128 bp each. Instead of aligning full-length inputs, we assessed leakage at the resolution of prediction, stratifying test set predictions for each 128 bp bin by its similarity to training set bins (maximum SW alignment score). (**B,C**) Model performance and memorization for (**B**) Median MSE and (**C**) Median Pearson correlation (*y*-axes) across all tracks for transcription factor binding ChIP, DNase, histone modifications ChIP, and CAGE assays (subplots), stratified by maximum SW alignment score (*x*-axes). (**D**) Scatterplot showing the relationship between model memorization (Pearson correlation between predictions and training sequence neighbor predictions, *x*-axis) and model performance (Pearson correlation between predictions and measurements, *y*-axis).

Similar performance-sequence similarity trends to those seen in other models were seen with Enformer. For all track types, the numerical prediction values appear to be anchored to those seen during model training, resulting in a MSE that decreased monotonically with increasing sequence similarity (**Fig. 6B**; **Supplementary Figure 14**). Pearson correlation between predictions and ground truth varied across homology ranges for all track types (**Fig. 6C**; **Supplementary Figure 15**). Homology effects varied across track types in ways consistent with each assay’s dependence on local sequence context. For transcription factor binding ChIP, performance was worst at intermediate-to-high homology (80-120 SW score) and high at both extremes of similarity (**Fig. 6C)**, as seen in MPRA models and ChromBPNet (**Fig. 3B**, **Fig. 5A**). Quantifying memorization as before (**Fig. 5B**) showed that memorization exceeded performance across the entire high-homology range, consistent with the model relying on memorization and being misled by similarity to training sequences when function diverged (**Fig. 6C)**. DNase showed a similar U-shaped relationship between sequence similarity and performance, but performance recovered more substantially as sequences became less similar to the training set, and memorization only briefly rivalled performance (90-100 SW) before falling (**Fig. 6C**). In both cases, the more memorization contributed, the worse the predictions. In contrast to TF ChIP and DNAse, where the activity observed is encoded in the sequence at that position, histone modifications typically arise from the activity of adjacent regulatory sequences. Accordingly, there was no performance boost due to memorization and instead performance increased monotonically as sequences became less similar to training data (**Fig. 6C**). CAGE, a measure of RNA polymerase II initiation, is also a product of the DNA sequences surrounding the bin containing the CAGE signal (e.g. enhancers, promoters) (76). CAGE showed consistently low memorization across all homology ranges and no clear relationship between performance and similarity to training sequences (**Fig. 6C**). Together, these patterns suggest that features that can be predicted locally (e.g. TF binding and chromatin accessibility) are especially susceptible to memorization compared with those that require integration of the broader sequence context (histone marks and CAGE). However, as noted below, memorized sequences can also impact predictions in adjacent bins, but would not be detected in this analysis.

We next considered how performance was impacted by memorization at the bin level. Considering all track and sample types, we found that memorization and performance were negatively correlated (Pearson *r* = −0.312; **Fig. 6D**). Furthermore, stratifying the tracks by their overall predictive success revealed that this detrimental impact is more pronounced for tracks where the model generally struggled to predict the variation across test bins (**Supplementary Figure 16**). It is possible that certain tracks are not easily explained by DNA sequence, driving memorization as the only way of decreasing the loss during training. However, this is unlikely to be the main explanation because performance is worse when the model has seen similar sequences during training compared to having never seen similar sequences before (**Fig. 6C**), supporting memorization driving the decreased performance at the bin level.

### Homology-biased performance at the locus level

While our bin-level analysis revealed signs of local memorization, the appeal of long-context models is that they can capture interactions between elements. Accordingly, we next asked how the total homology across entire input sequences impacted Enformer predictions. We summarized each test sequence by calculating the median of its bin-level maximum SW alignment scores (**Fig. 7A**), and then, for each sequence and each track, we computed the Pearson *r* between predicted and observed profiles across all 896 output bins (30,31). Using H3K9me3 ChIP-seq as an example, the extremes of homology to training data reveal stark performance differences (**Fig. 7B**); high overall similarity (median homology score = 112) exhibited poor correspondence between predictions and experimentally determined values (*r* = 0.202) and low similarity (median homology score = 0) had a good correspondence (*r* = 0.648). When we scaled the analysis to all test sequences and assay types, we observed a clear and consistent inverse relationship between homology between training and test data and model performance (**Fig. 7C**), with the highest performance again concentrated in the lowest homology ranges. This trend was consistent across the vast majority of tracks, including CAGE (**Fig. 7C**). This is consistent with sequences similar to training data having their fine-scale regulatory variation obscured by memorization, while sequences with little training-set similarity force the model to rely on learned regulatory logic that better captures the output profile.

**Figure 7:**
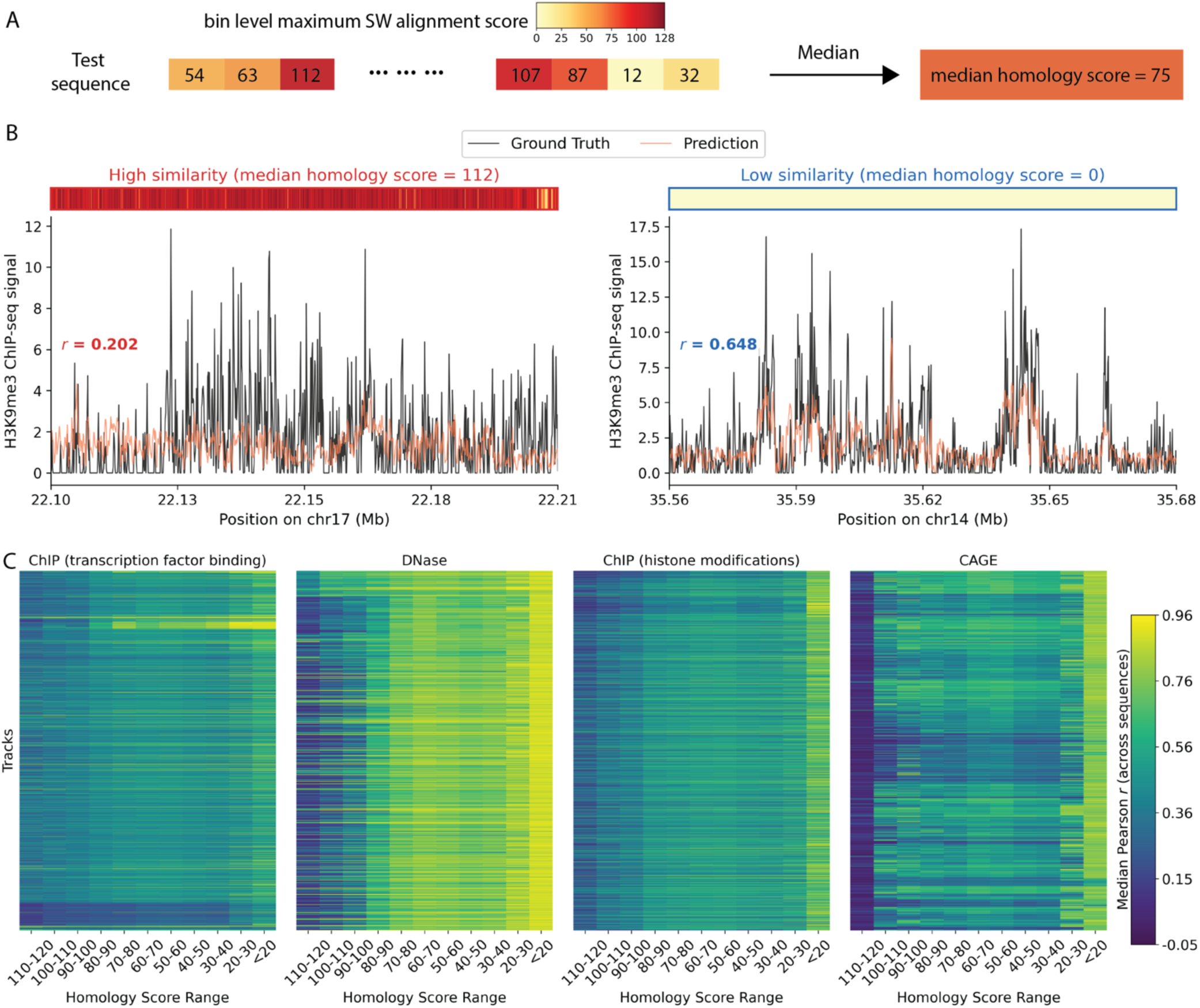
Overall homology systematically biases Enformer predictions across sequences. (**A**) Schematic of locus-level homology quantification. For each test sequence, a homology score is calculated as the median bin-level maximum SW alignment scores (colored by magnitude) across all output bins. (**B**) Example Enformer predictions for H3K9me3 ChIP-seq for test sequences with the highest median homology score (left) and lowest median homology score (right). Heatmaps above each plot show the bin-level maximum SW alignment scores across 896 bins (*x*-axis). Ground truth (black) and model predicted (red) H3K9me3 ChIP-seq signal (*y*-axes) are shown across bin positions along the sequence (*x*-axes). (**C**) Heatmaps of per-track median Pearson correlations (color scale) across sequences within each homology score group (*x*-axes) for all tracks (*y*-axes) across transcription factor binding ChIP (1993 tracks), DNase (674 tracks), histone modifications ChIP (1998 tracks), and CAGE (638 tracks) (subplots).

## Discussion

In this study, we demonstrated that homology confounds the training of sequence-to-function models across all tested modeling regimes, spanning simulations, fixed-context MPRA models, and short and long context genome-trained models. A model that generalizes well should have no relationship between how similar test sequences are to its training data and its ability to predict their function, but for all models we saw the opposite (**Fig. 3B, 5A, 6E, 7C**): models generally perform well on sequences that are unrelated to those observed during model training but have variable performance on homologs to training sequences according to the model and the degree of homolog similarity. High performance on non-homologous sequences suggests that models have indeed learned generalizable regulatory features. While high performance on close homologs appears at first to suggest we should train our models on close homologs of the sequences we care most about, our results suggest that exactly the opposite is true. Instead, the high performance when training on close homologs appears to result from non-causal associations being learned by the models. This is illustrated by all models performing worse when training on certain homologs (close homologs for Enformer, distant homologs for others) compared with training on no homologs at all. Drawing a parallel from DNA language modeling, down-weighting repetitive sequences has been shown to improve model performance (59), consistent with the idea that homologous sequences encourage memorization at the expense of learning generalizable sequence features.

The performance profiles across modeling regimes further illustrate when memorization pays off and when it backfires. In MPRA (**Fig. 3B**) and ChromBPNet (**Fig. 5A**), performance consistently peaked at the highest homology ranges, but for Enformer this trend was inverted. In MPRA, sequences are assayed in an identical reporter, so two similar sequences share both composition and functional context, and memorized associations tend to transfer. For ChromBPNet, the prediction window is small enough that similar sequences tend to share the regulatory context driving accessibility, so memorized associations still largely transfer. For Enformer, however, two bins that are locally similar at 128 bp sit in entirely different ∼196 kb contexts, so the broader regulatory environment that ultimately determines each bin’s activity will generally differ. Memorization therefore anchors predictions to training neighbors whose function is unlikely to be conserved, and performance at the most homologous bins falls below what the model achieves at low homology score ranges when forced to rely on generalizable regulatory logic. Consistent with this, the strong performance at high similarity in MPRA and ChromBPNet appears to be partly a product of memorization as well.

Beyond these memorization effects, homology may also compromise the training data itself. Reads originating from highly similar loci can map ambiguously, causing signal from one locus to be attributed to a related locus elsewhere in the genome (77–79). This would cause homologous regions to share similar assay readouts not only because of shared function but also as an artifact of read misassignment, further reinforcing the association between sequence similarity and shared signal that models can exploit. While we do not directly quantify this effect in our study, it represents an additional mechanism by which homology can compromise both model training and evaluation, and warrants further investigation.

These findings have direct implications for variant effect prediction. The prevalence of SNV doppelgängers (**Fig. 2D, 2E; Supplementary Figure 5**), where alternate alleles with flanking sequences are replicated elsewhere in the genome, creates fundamental ambiguity in model evaluation: when a model predicts a variant’s effect, it remains unclear whether it has learned cis-regulatory logic or memorized a sequence-function relationship seen during training. Homology with the training data could lead the SNV to be correctly predicted because the answer is the same as in the training data, or incorrectly predicted because the model was misled by the homology in the training data. Consequently, it will be important to identify whether models used for predicting the mechanisms underlying variant effects have seen homologous sequences in the training data, and to account for this when interpreting model predictions. Given the abundance of SNV doppelgängers, this issue is likely to confound current estimates of variant effect prediction model performance. hashFrag can address this by identifying cases where sequences of interest were seen during model training, enabling homology-aware stratification and reliable interpretation of variant effect predictions.

Beyond evaluation, we demonstrated that hashFrag-trained models outperform chromosomally split models when evaluated on synthetic sequences with no genomic homology (**Fig. 4G, Supplementary Figure 11**), indicating that chromosomal splitting creates inferior train-validation splits that lead to suboptimal model training and selection. There are two likely reasons for this. First, because hashFrag permits the utilization of sequences from all chromosomes during training it may produce more functionally diverse training examples (e.g. X chromosome inactivation and escape occur only on X (80)). Second, by eliminating leakage between training and validation, hashFrag can provide a more accurate estimate of overfitting (**Fig. 1C**), leading to better model selection. Based on these results, for most applications we advise strict removal of homology-based leakage for validation sets to ensure the selection of generalizable checkpoints. Counter-intuitively, hashFrag should generally not be used to construct fully orthogonal train-test splits. Homology is a structural feature of the genome, so it inevitably permeates the training data regardless of how splits are constructed. Models would still have homology-based biases under orthogonal splits, but these biases would go undetected because the test set would contain no homologous sequences to expose them. We recommend stratification of test data by homology to training sequences to evaluate generalization across sequence similarity ranges.

The speed and sensitivity of hashFrag make it extremely useful in characterizing homology-based leakage even in large sequence datasets. Selecting a specific threshold for determining homology is challenging as there is no definitive alignment score at which sequences transition from homologous to non-homologous. A threshold that is too strict risks eliminating convergently evolved sequences (e.g. CTCF binding sites) from either training or validation data, thereby hindering the model’s ability to learn some aspects of sequence function. In contrast, a threshold that is too lenient may include some homologous sequences. Accordingly, we recommend dinucleotide shuffling a subset of sequences and selecting a threshold larger than the largest score seen for the dinucleotide-shuffled sequences (e.g. ∼60 for 200 bp genomic sequences; **Fig. 2B, Supplementary Figure 2**). While this approach will undoubtedly include some very distant homologs, based on our results, distant homologs differ substantially in their activity and are especially difficult for the models to predict (**Fig 3B, 5A**), and so generalization estimates will not to be overestimated at a reasonably lenient threshold choice. hashFrag can also be used to detect inter-species leakage during training while augmenting data with sequences from other organisms (81–84), and identifying cases where sequences of interest were used during model training when performing downstream inference tasks (e.g. variant effect prediction).

Longer sequence models provide a unique challenge for leakage analysis. In addition to being computationally intensive to align, the longer the sequence, the more likely two sequences are to share local fragments with at least some homology, making truly orthogonal splits difficult if not impossible (27). While we demonstrated how homology affects predictions locally (using the 128 bp bins of Enformer; **Fig. 6**), this approach cannot fully account for the influence of homologous fragments elsewhere in the ∼196 kb input window. Indeed, our analysis revealed that more sequence-intrinsic features have clearer relationships between sequence similarity to training data and performance (e.g. MPRA, open chromatin, transcription factor binding ChIP; **Fig. 3B, 5A, 6E**), whereas products of distal regulation (e.g. gene expression via CAGE) have more complex relationships (**Fig. 6C**). Indeed, while we saw that Enformer’s performance was negatively affected by leakage at the locus level (**Fig. 7**), it is clear that in some cases leakage could also result in correct predictions for wrong reasons. For example, a non-leaked transcript could be correctly predicted because the model has memorized the activity of a nearby leaked enhancer. While one could mask the predictions at the leaked regions during model evaluation, distal regulation allows leaked sequences to influence predictions of neighboring non-leaked sequences, and so the potential for leakage in evaluation remains. Fully characterizing distal leakage remains an open problem, and additional metrics accounting for long-range interactions will need to be developed.

Our findings suggest that the consequences of homology-based leakage depend not only on genome structure itself, but also on which models are used, how they are trained, and what they are trained on. Our simulation revealed that model capacity can influence the degree of memorization (**Fig. 1**), and showed that DREAM-RNN and ChromBPNet differed in their memorization despite being trained on the same data (**Fig. 5C**). Interestingly, DREAM-RNN was optimized on sequence-expression data of random DNA sequences (66), which are ideal for optimizing models to be maximally generalizable because every sequence is singular in the training data, creating a balanced dataset, and random DNA has no homology structure, allowing generalization to be estimated robustly with the independent validation data. Homology-based leakage appeared to be especially detrimental to Enformer (**Fig. 6, 7**), suggesting that preventing memorization would improve its performance. Looking forward, it will be important to explore training objectives that explicitly account for homology structure in genomics data, for example by reweighting homologous examples to correct the effective training data imbalance caused by our repetitive genomes. We also provide an approach for quantifying memorization in genomics models. Importantly, our approach does not require that the sequence have homologs in the data, nor does it even require the sequences to actually have been measured – it gauges similarity between predictions on training sequences to predictions of related sequences. Quantifying how quickly predictions change as training sequences diverge thus provides a straightforward way of quantifying memorization, with the objective of matching the experimentally determined divergence rate (85,86). Finally, reporting homology-stratified metrics, ideally using consistent data splits, will be critical in monitoring progress. Genomic sequences violate one of the central tenets of machine learning: that data are independent; recognizing and accounting for this will be essential for building predictors that learn truly causal models of genome function.

## Methods

### Simulation of homology-based model overestimation

We performed a controlled simulation to isolate the effects of homology-based data leakage on model evaluation. 1,000 random 200 bp DNA seed sequences were generated composed of uniformly sampled nucleotides. Half of the sequences were designated as the positive set and had 1–3 transcription factor binding sites (TFBSs) randomly inserted from a curated set of 20 TFBSs (10 bp length); the remaining half served as the negative set with no TFBSs embedded. For each seed sequence, 20 homologous variants were generated by introducing random point mutations at a 5% rate across all positions for negative sequences, and at a 5% rate restricted to non-TFBS regions for positive sequences to preserve the regulatory signal. This yielded a final population of 20,000 sequences with defined homology structure, where the true discriminative feature between classes was the presence or absence of TFBSs.

80% of origins were randomly selected for training-associated data and, from each such origin, sampled 8 homologs for the training set (yielding 6,400 sequences, then randomly subsampled to 6,000). The remaining homologs from these same training origins formed a pool from which we sampled 2,000 sequences to create a leaky hold-out set. To create a homology-aware set, the remaining 20% of origins were used (disjoint from training origins) and 2,000 sequences were sampled, guaranteeing that no sequence shares an origin with any training sequence.

We trained convolutional neural networks (CNNs) of three capacity levels to predict sequence class from one-hot encoded sequences. The Shallow CNN consisted of a single convolutional layer (16 filters, kernel size 11) followed by batch normalization, ReLU activation, adaptive average pooling, and a linear output layer. The Medium CNN added a second convolutional layer (32 and 64 filters, kernel sizes 11 and 7, respectively) with max pooling between layers. The High-Capacity CNN comprised three convolutional layers (64, 128, and 256 filters; kernel sizes 15, 7, and 7) with batch normalization after each layer, max pooling after the second and third layers, dropout (p=0.5), and a two-layer classifier head (256→128→1) with an additional dropout layer. All models used sigmoid activation for binary output. Models were trained using the Adam optimizer with a learning rate of 0.01 and binary cross-entropy loss for 100 epochs. Model performance was evaluated using area under the precision-recall curve (AUPRC).

### MPRA Dataset and model training

We used a large-scale MPRA library measured in lymphoblasts (39). The library comprised approximately 226,000 sequences in both forward and reverse orientations, derived from promoters and putative enhancers specific to K562 cells. The dataset was split chromosomally, designating around 60,000 sequences for training and the remaining sequences for testing. Three distinct chromosomal splits were performed as outlined below:

Split 1: Training set chromosomes 6, 2, 4, 21, 9, 18, 13

Split 2: Training set chromosomes 16, 14, 21, 7, 12, 22, 5

Split 3: Training set chromosomes 17, 6, 18, 10, 22, 13, 11

In all the splits above, held out sequences are used as the test set. It should be noted that the number of chromosomes in the training sets were limited to retain a larger portion for testing, enabling finer-resolution stratification by alignment score.

DREAM-RNN, DREAM-CNN, DREAM-Attn, and MPRAnn were trained using the chromosomal splits following the training specifics as described in respective models from (66) and (39). Model performance is evaluated using Pearson correlation on the different subsets of the test set, with sequences separated by their maximum SW local alignment to any sequence in the training set.

### SW Local Alignment Calculation

To quantify sequence similarity, we employed the SW local alignment algorithm (54), which uses dynamic programming to align subsequences and compute alignment scores based on match, mismatch, and gap penalties. Match score 1, mismatch score -1, gap open score -2, and gap extension score -1 were selected to compute the alignment score using the Biopython implementation (87).

### Maximum Pairwise SW Local Alignment

For each sequence in a set, the SW local alignment scores were calculated against all sequences in the other set and the maximum score was taken. This is likely to underestimate the degree of leakage as some sequences are likely to have different parts match different training sequences, and it also does not account for the number of matches in the training data.

### OverfitNN

OverfitNN, short for Overfit Nearest Neighbor, is a heuristic tool designed to predict gene expression purely based on memorization, without utilizing any cis-regulatory logic. It is a nearest-neighbor approach, where for a given test sequence, it queries the training set to identify the top-k closest sequences within a range of SW alignment scores. Prediction is made by calculating the average expression of these nearest neighbors. In our analysis, OverfitNN was applied to different chromosomal train-test splits, with the nearest neighbor set to 10 (**Fig. 3A**, **Supplementary Figure 6**).

### SNV Doppelgänger

GWAS SNVs with PIPs greater than 0.1 were selected from the OpenTargets dataset (56), yielding a set of variants likely to be causally associated with some trait or condition. To identify SNV doppelgängers, that is, occurrences of the alternate allele along with its surrounding sequence context elsewhere in the genome, the DNA sequence at each SNV was extracted from the reference genome, the alternate allele inserted, and the resulting sequence used as a query to find exact matches elsewhere in the genome.

### hashFrag

hashFrag can filter homology-based leakage from test data for existing train-test splits. It can be used to stratify the test dataset based on the degree of homology-based leakage. Moreover, it can divide a sequence dataset into leakage free training, validation, and test splits.

hashFrag uses BLASTn which is a heuristic algorithm that can be used to identify local regions of similarity between DNA sequences. A BLAST database composed of all sequences in the dataset is constructed, and then each sequence is queried against this database to identify candidate sequences it exhibits similarity to. This gives a list of candidate homolog pairs along with their heuristic BLAST alignment score. hashFrag has two modes, hashFrag-lightning and hashFrag-pure. In hashFrag-pure, the exact SW score is calculated using biopython due to BLAST scores being heuristic, whereas BLAST scores are used directly in hashFrag-lightning. Calculating precise SW score is optional as it increases the runtime by about 2x (**Fig. 4C**) and the homologs missed are near the threshold anyhow (**Supplementary Figure 8**), and so unlikely to result in substantial leakage.

After identifying pairs of sequences exhibiting sufficient similarity to be considered homologous, homology-aware data splits are created using a graph-based approach. By representing each sequence as a node and homology between two sequences as an undirected edge, groups of homologous sequences are readily identified. Specifically, different disconnected subgraphs represent different clusters of sequence homology, and isolated sequences with no edges represent “orthogonal” sequences with no homology to any other sequence in the set. Data splits can then be created by sampling connected subgraphs into each split, effectively ensuring that no homologous sequences end up in different splits.

### Discrepancy between alignment scores of BLASTn and biopython

There are two reasons BLASTn scores and biopython SW scores can differ. Firstly, BLAST uses heuristics and may miss the optimal alignment. Secondly, even for the same alignment BLASTn and biopython may report different scores due to different scoring methods. Namely, while penalizing gap extensions, BLASTn penalizes the total number of gaps (applying penalties for both gap openings and gap extensions), although gap openings are penalized separately, leading to a double counting effect. To ensure comparability with exact SW scores calculated using biopython, this double counting of gap openings was corrected in the BLASTn scores, aligning the scoring mechanism with that of Biopython, where gap openings and extensions are treated separately. Additionally, it is worth noting that reducing the word size in BLASTn can yield alignments closer to the optimal alignment at the cost of increased runtime.

### Comparison of hashFrag-pure and hashFrag-lightning

In the hashFrag-pure mode, BLAST is used to generate candidates, which are later filtered based on their exact SW local alignment scores, ensuring homology is assessed against a specified threshold. This filtering step is needed because BLAST scores, while a strong approximation, may not provide exact SW alignment scores. For detailed analyses on test datasets, we recommend hashFrag-pure as the false-positive removal step minimizes leakage, as only sequences missed by BLAST will contribute to residual leakage. However, if the primary objective is only to create train-test splits, hashFrag-lightning can be used where BLAST scores alone are used without additional precise SW calculation, offering a faster alternative while maintaining reasonable accuracy (**Supplementary Figure 8**).

### Tuning BLAST parameters

We initially tuned BLASTn parameters on subsamples of 10,000 sequences from the MPRA dataset (**Supplementary Figure 17**) to maximize its ability to capture possible instances of sequence homology (i.e., maximize recall capabilities). The following parameters were screened: the length of exact sequence match required to initiate alignment, *word_size* (5, 7, 11, 16 and 20); the maximum number of target sequences that can be returned for a given query sequence, *max_target_seqs* (500, 1000, 5000 and 10000 (all)); the statistical significance of the alignment, or Expect value (E-value), threshold to return an alignment, *evalue* (10, 50 and 100), and whether to mask homology stemming from low-complexity regions such as repeats, *dust* (“yes” or “no”).

A grid-search was performed over all possible combinations of the parameters on the subsampled datasets. Evaluating the recall and FPR of homologous sequences in terms of the SW scores, a word_size of 7, max_target_seqs set to the total number of sequences in the dataset, an evalue of 100 and no dust filter were selected as parameters for BLASTn on the complete MRPA dataset. After determining candidate groups of similar sequences for every sequence in the dataset, false-positive collisions were filtered according to a SW threshold of 60 (**Supplementary Figure 18**), which was chosen based on the range of SW scores observed in dinucleotide shuffled sequences.

### hashFrag performance analyses

Given all pairwise SW local alignment scores for sequences, we defined thresholds spanning the possible range of observed SW scores (i.e., [0,200]) to define the score above which the match is considered a positive example of homology. According to each threshold, true-positive, true-negative, false-positive, and false-negative statistics were calculated for all pairs of sequences with the positive set defined by whether the pair met the SW score threshold and the predicted positives corresponding to whether the pair was detected by BLASTn. This allowed further calculation of recall and false-positive rate statistics (**Fig. 4B**). In terms of evaluation, recall was prioritized as a metric based on the rationale that false-negatives can directly manifest as leakage in the eventual data splits, whereas false-positives have no effect on leakage.

### hashFrag train-test leakage removal/test set stratification

A BLAST database composed of all sequences in the training dataset is constructed, and then each test sequence is queried against this database to identify candidate sequences it exhibits similarity to. Exact SW score is calculated for the BLAST candidates then sequences are removed from the test split based on a threshold or stratified based on ranges of SW scores.

### Computing costs

The high cost of computing many pairwise sequence comparisons is one of the main obstacles that users must confront in order to mitigate data leakage due to sequence homology in train-test splits. To that end, CPU runtime of the executed high-performance computing (HPC) jobs was used to characterize the time cost of performing such an analysis. To maintain memory cost at 16 gigabytes and a single computing node, pairwise computations were partitioned into smaller batch jobs and later aggregated via summation to obtain the final CPU time cost. The naive method of computing all pairwise comparisons between sequences in the dataset was compared against the procedure of performing hashFrag to identify candidate pairings and then subsequently computing pairwise SW scores only among those candidates.

### Comparison of homology between hashFrag *vs*. chromosomal splits

Using hashFrag-pure, homology-aware data splits were created (80% train; 20% test) at a local alignment threshold of 60. A chromosomal train-test split was also created following the same ratio. The maximum pairwise local alignment scores between train and test sets were then used to quantify the degree of similarity (**Fig. 4D**).

### Mitigation of data leakage

We removed sequences in the test split that exhibited high SW scores with sequences in the train split, applying different SW score thresholds. Pearson correlation coefficients were calculated based on the available data points after filtering at each specific threshold (**Fig. 4F**; **Supplementary Figure 9**).

### Comparison between hashFrag vs chromosomal split trained models

We trained 100 replicates of DREAM-CNN, DREAM-RNN, and DREAM-Attn models on the Gosai et al. (13) genome-derived MPRA dataset using different training and validation splits generated by both hashFrag and chromosomal splitting methods, downsampling to ensure that all splits contained the same number of sequences (160,000 for training and 40,000 for validation). Chromosomal splits were created by sending chromosomes to training and validation sets until the total number of sequences exceeded 160,000/40,000 and then downsampled to make the number of sequences exactly 160,000/40,000. hashFrag splits were created by splitting the dataset in 80-20 ratio and then downsampling to make the number of sequences exactly 160,000/40,000. The best model was selected based on Pearson correlation on the validation data. We used 17,000 designed sequences from Gosai et al. (13) as the test data (**Fig. 4G**, **Supplementary Figure 11**), which are very dissimilar to the genomic sequences (**Supplementary Figure 10**).

### Scaling for large sequence datasets using hashFrag HPC module

To enable homology analysis of genome-trained models with large training sets and longer sequences, we extended hashFrag with a distributed computing module for execution on HPC clusters. The distributed hashFrag module partitions the analysis into independent jobs suitable for array job submission. First, a BLAST database is constructed over all sequences in the dataset. The query sequences are then partitioned into chunks based on a user-specified partition size. Each chunk is configured as an independent array job that queries its sequences against the complete database. The module generates submission scripts compatible with SLURM and Sun Grid Engine (SGE) job schedulers.

### Short-context genomic model task

We used ChromBPNet (33) and chromatin-specific DREAM-RNN (66) models trained on K562 ATAC-seq data. Both models were trained as described in their respective original publications (33,66). hashFrag was applied with the HPC module to compute pairwise SW alignment scores between all test and training sequences, and test sequences were than stratified by their maximum alignment score to the training set for performance evaluation.

### Model memorization

To quantify the extent to which model predictions reflect memorization of training data, we developed a memorization metric. For each test sequence, the most similar training sequences were identified based on maximum SW alignment scores and computed the memorization prediction as the mean of the model’s predictions on these similar training sequences. The Pearson correlation or mean squared error (MSE) were calculated between the model’s test predictions and the corresponding memorization predictions, where lower MSE and higher Pearson correlation indicate higher memorization.

### Enformer bin-level analysis

Enformer takes 196,608 bp input sequences and predicts regulatory track values across a center 114,688 bp region divided into 896 bins of 128 bp each. The pretrained Enformer model and its training and test sequence coordinates were obtained from the original publication. For each of the 1,937 test sequences, 896 bins (128 bp each) comprising the center prediction region were extracted. This yielded approximately 1.7 million test bins for analysis. For each test bin, maximum SW local alignment scores were calculated against all bins from all training sequences using the hashFrag HPC module. Test bins were stratified into groups based on their score ranges (e.g., 20-30, 30-40, …, 120-128). For each score range, model performance and memorization were computed using Enformer’s predictions on the 1,937 test sequences and the ground truth experimental values across all 5,313 human tracks.

### Summarizing bin-level performance across Enformer output tracks

Enformer predicts 5,313 human regulatory tracks simultaneously for each input sequence. To summarize model performance and memorization within each homology score group, the relevant metric (Pearson *r* or MSE) was calculated independently for each track and then the median across all tracks was taken. This median-of-tracks approach provides a summary of model behavior that is not driven by outlier tracks with unusually high or low values. When examining track-type-specific patterns, the same procedure was applied within each assay class (ChIP, n = 3,991 (transcription factor binding ChIP, n = 1993 and histone modifications ChIP, n = 1998); DNase, n = 674; CAGE, n = 638), taking the median across tracks of the same type.

### Enformer locus-level homology

For the locus-level evaluation, a similarity score was calculated for each test bin, but aligned it against full 114 kb training sequences (center of 196 kb) rather than 128 bp training bins. For each 128 bp bin within a test sequence, the maximum SW local alignment score was computed between that bin and any full training sequence. This adjustment is necessary because a homologous match can fall anywhere within a training sequence, including across the boundary between adjacent bins; bin-versus-bin alignment would split such matches and systematically underestimate similarity for partially overlapping homologs. Each test sequence was summarized by the median of its 896 bin-level scores, which captures the typical homology burden of the sequence while minimizing the influence of individual outlier bins. The 1,937 test sequences were stratified by their median homology scores, and for each stratum we computed locus-level performance as the Pearson *r* between predicted and observed profiles across all 896 bins, separately for every track.

### Enformer locus-level performance and track aggregation

We evaluated Enformer’s locus-level performance by calculating the Pearson correlation between the predicted and observed profiles across all 896 bins for every individual track. To assess the generalizability of homology–performance trends across individual regulatory targets, the mean locus-level Pearson correlation was computed for each track independently, and averaged across all test sequences falling within a given locus-level homology score group (**Fig. 7C**).

## Data availability

The processed datasets used for this study are available from Zenodo (https://doi.org/10.5281/zenodo.14715096).

## Code availability

Open-source code for our tool is available from GitHub (https://github.com/de-Boer-Lab/hashFrag). Additionally, the code used for the analyses presented in the paper can be accessed here (https://github.com/de-Boer-Lab/hashFrag-PaperCode).

## Acknowledgements

We are grateful to Mateusz Faltyn and Grzegorz Boratyn for the valuable discussions. We are also grateful to the members of the de Boer Lab for their support during this work, including the cheerleading and sticker advocacy. This research was supported by the Stem Cell Network (ECR-C4R1-7), and the Canadian Institute for Health Research (PJT-180537). A.M.R. was supported by a UBC 4-Year Fellowship. C.G.D. is a Michael Smith Health Research BC Scholar. N.Y. was supported by Canada Foundation for Innovation (CFI)/B.C. Knowledge Development Fund (BCKDF), Canadian Institutes of Health Research (CIHR) Project Grant, Canadian Institutes of Health Research (CIHR) Canada Research Chair in Synthetic Biology, Allen Distinguished Investigator Award, and Canadian Institute for Advanced Research MacMillan Multiscale Human program. This research was enabled in part by support provided by the Digital Research Alliance of Canada, and Advanced Research Computing at the University of British Columbia.

## Author contributions

A.M.R. and C.G.D. conceived the study. A.M.R., C.G.D., and. B.K. designed experiments. B.K. developed the pairwise alignment scoring pipeline and the hashFrag codebase. A.M.R. trained models, developed methods to perform homology analyses, and evaluated model performance. A.M.R., C.G.D., and B.K. interpreted results. A.M.R., C.G.D., and B.K. wrote the manuscript. C.G.D. and N.Y. edited the manuscript. C.G.D. supervised the study. C.G.D. and N.Y. provided funding.

---

**Supplementary Figure 1:**
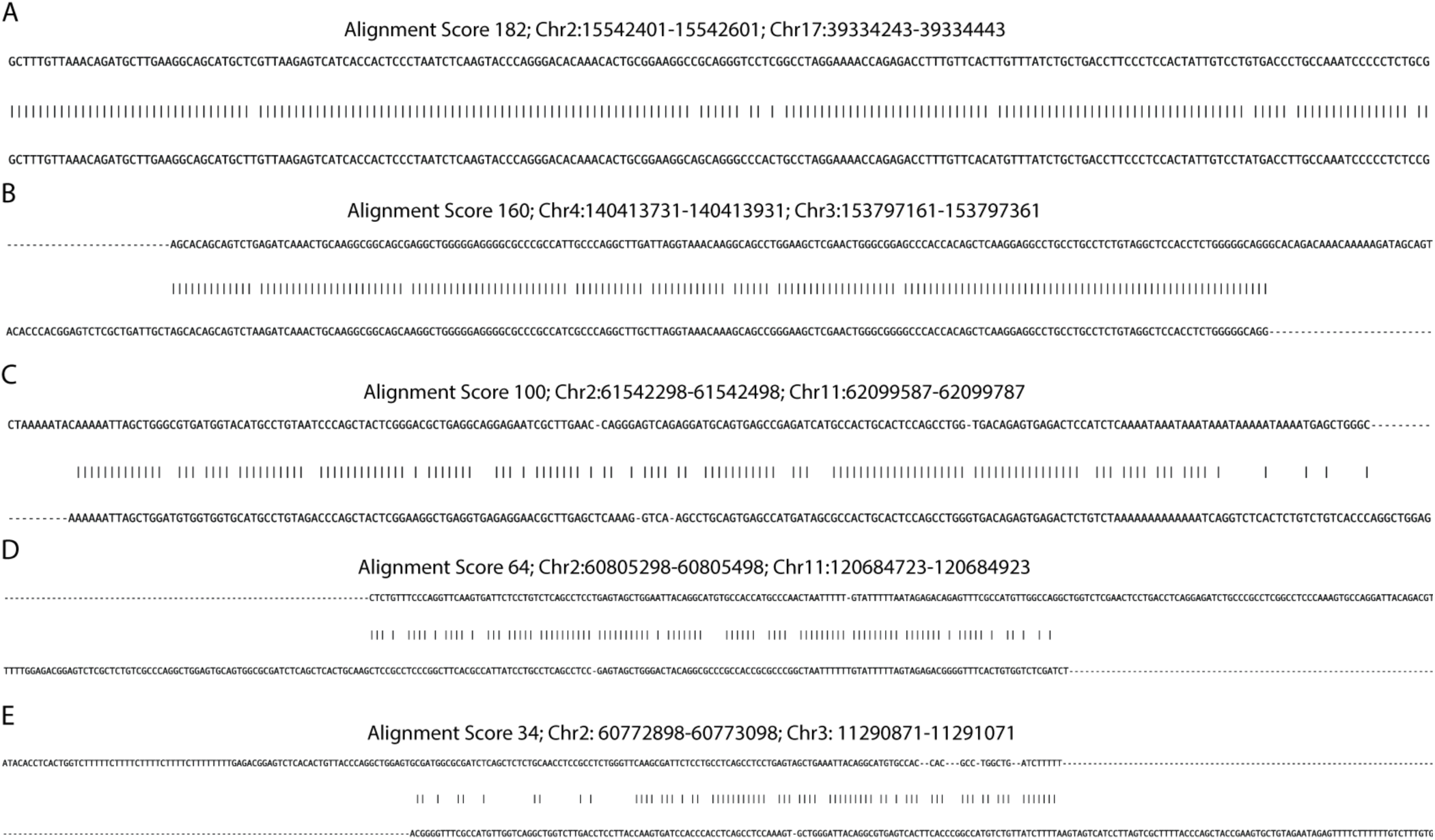
Examples of genomic sequences from different chromosomes showing different SW alignment scores. (A–E) Five sequence pairs from the MPRA dataset (39) are shown with varying degrees of similarity, indicated by alignment scores (A) 182 (chr2 *vs.* chr17), (B) 160 (chr4 *vs.* chr3), (C) 100 (chr2 *vs.* chr11), (D) 64 (chr2 *vs.* chr11), and (E) 34 (chr2 *vs*. chr3).

**Supplementary Figure 2:**
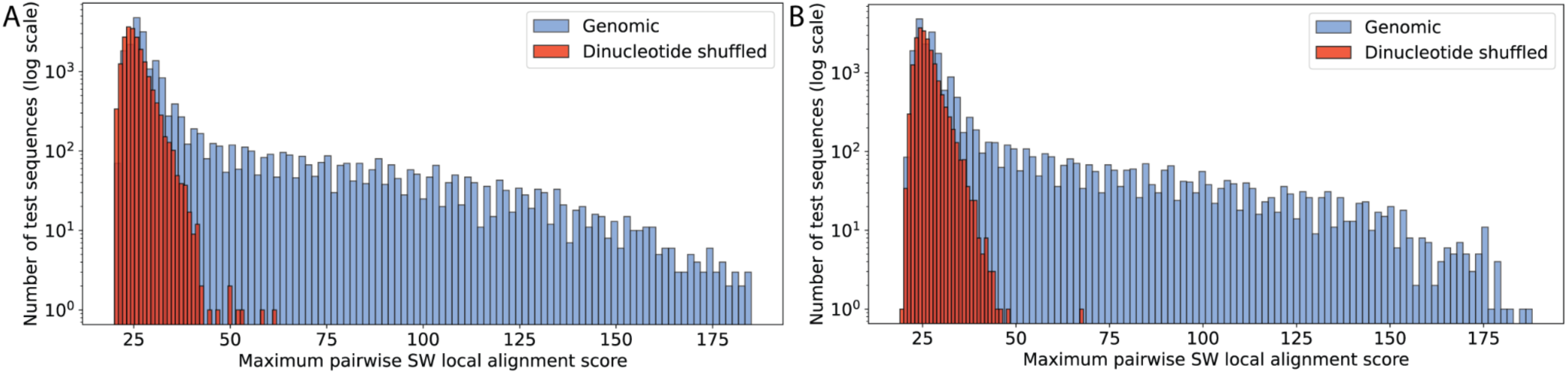
Genomic sequences show homology across chromosomes. **(A, B)** Histogram showing the number of test sequences (*y*-axes) with corresponding maximum pairwise SW local alignment scores with the training sequences (*x*-axes) for both genomic (blue) and dinucleotide shuffled (red) sequences, with training and test sets randomly sampled from distinct chromosome sets (20,000 each) for two different chromosomal splits (**A** and **B**).

**Supplementary Figure 3:**
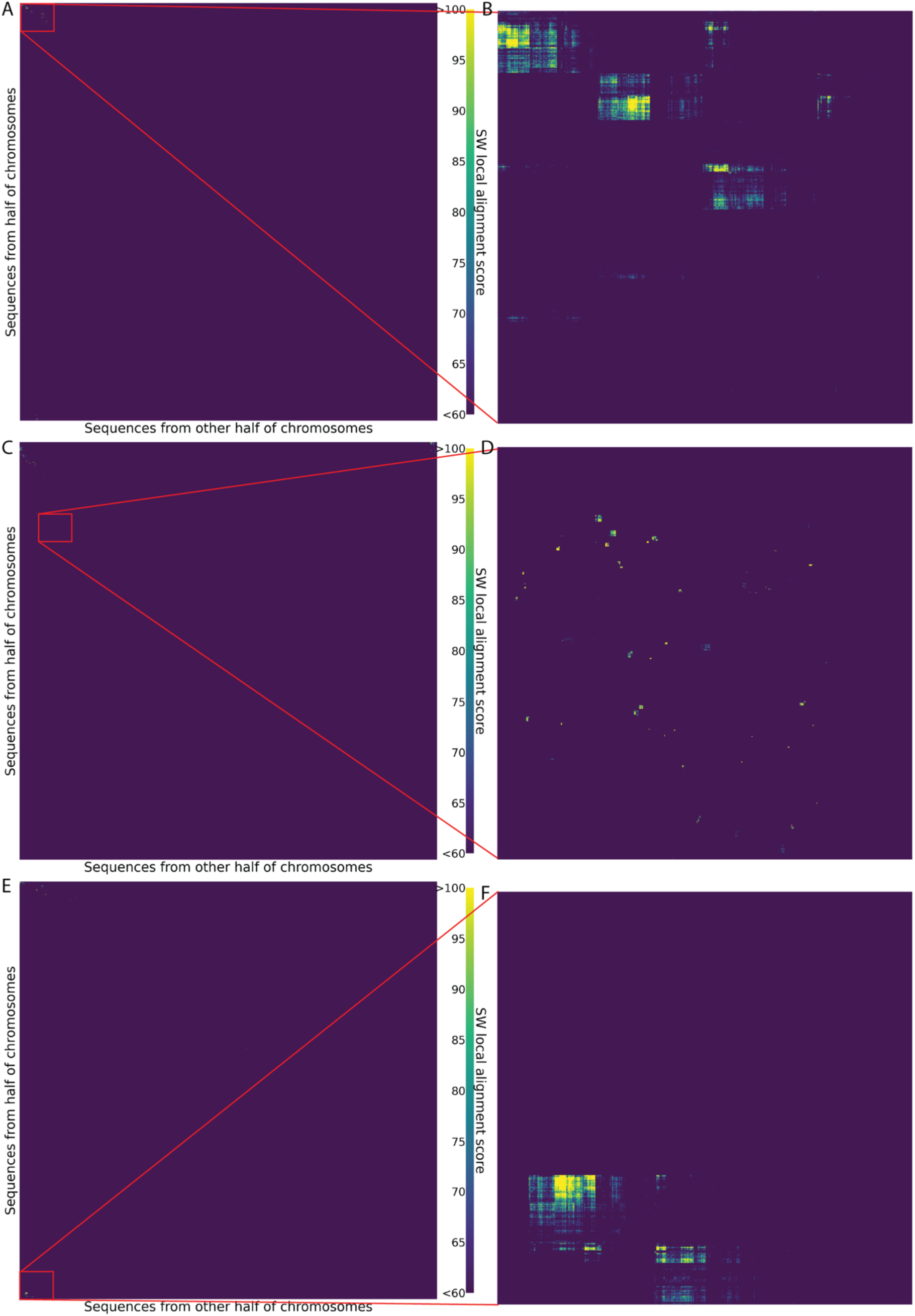
Genomic sequences show distinct clusters of homology. **(A, C, E)** Hierarchical clustering heatmap of local alignment scores between 20,000 sequences from one half of the chromosomes (*x*-axes) and 20,000 from the other half (*y*-axes) in chromosomal splits 1,2, and 3 (**Methods**). The heatmap range capped at alignment scores of 60 and 100, for ease of visualization, highlighting regions of homology vs. no homology. **(B, D, F)** Zoomed-in (1000 × 1000) view of the hierarchical clustering heatmaps.

**Supplementary Figure 4:**
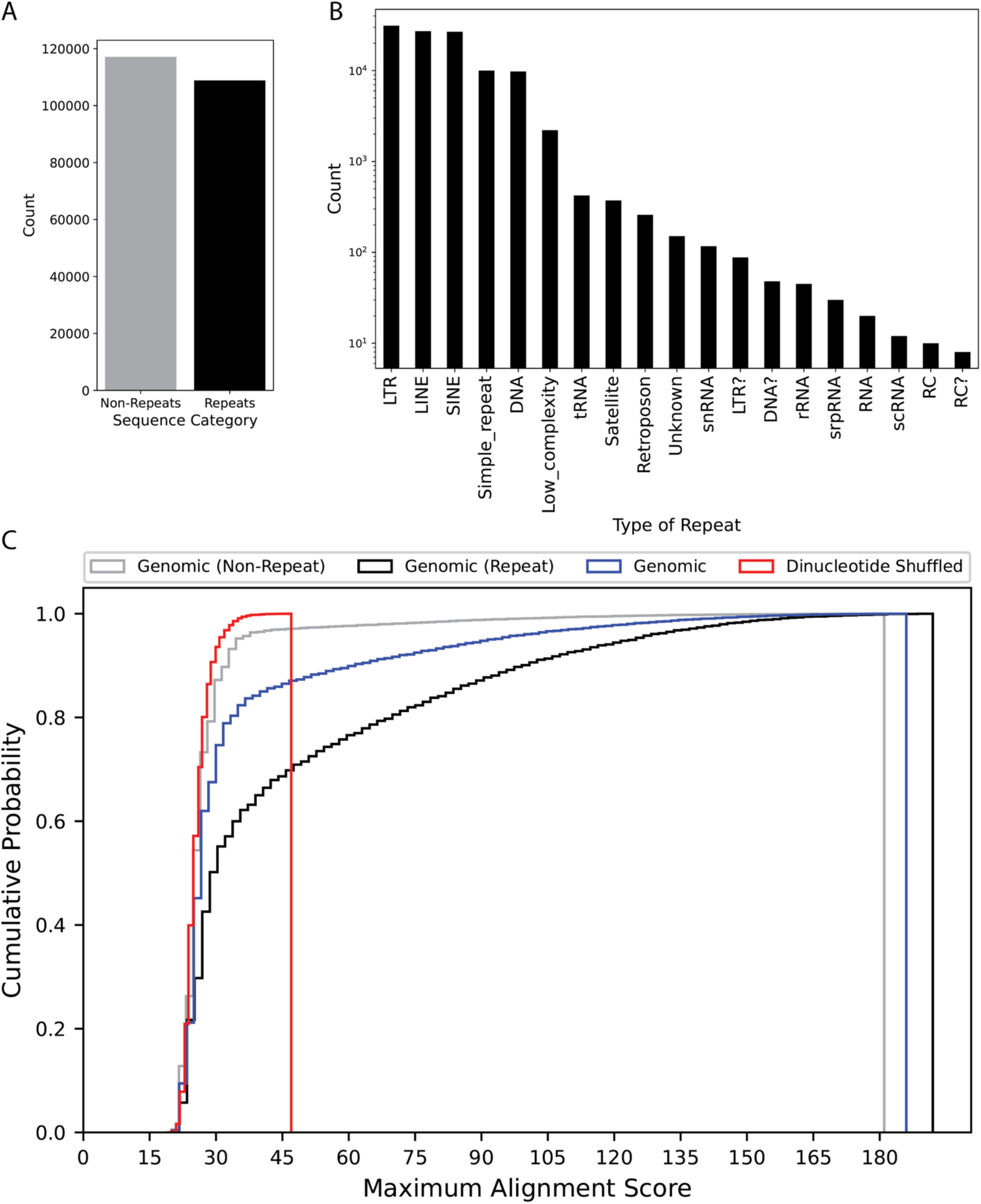
The majority of high-similarity sequences between chromosomal sets are from repeat elements. (**A**) The number of sequences (*y*-axis) from the MPRA dataset (39) that overlap or do not overlap repeat elements (*x*-axis). (**B**) The number of MPRA sequences (*y*-axis) that overlap each type of repeat element (*x*-axis). (**C**) The cumulative distribution (*y*-axis) of maximum alignment scores (*x*-axis) calculated between two chromosomally-split sets for non-repeat genomic sequences, repeat genomic sequences, all genomic sequences, and dinucleotide shuffled sequences (colors) in the MPRA dataset (39).

**Supplementary Figure 5:**
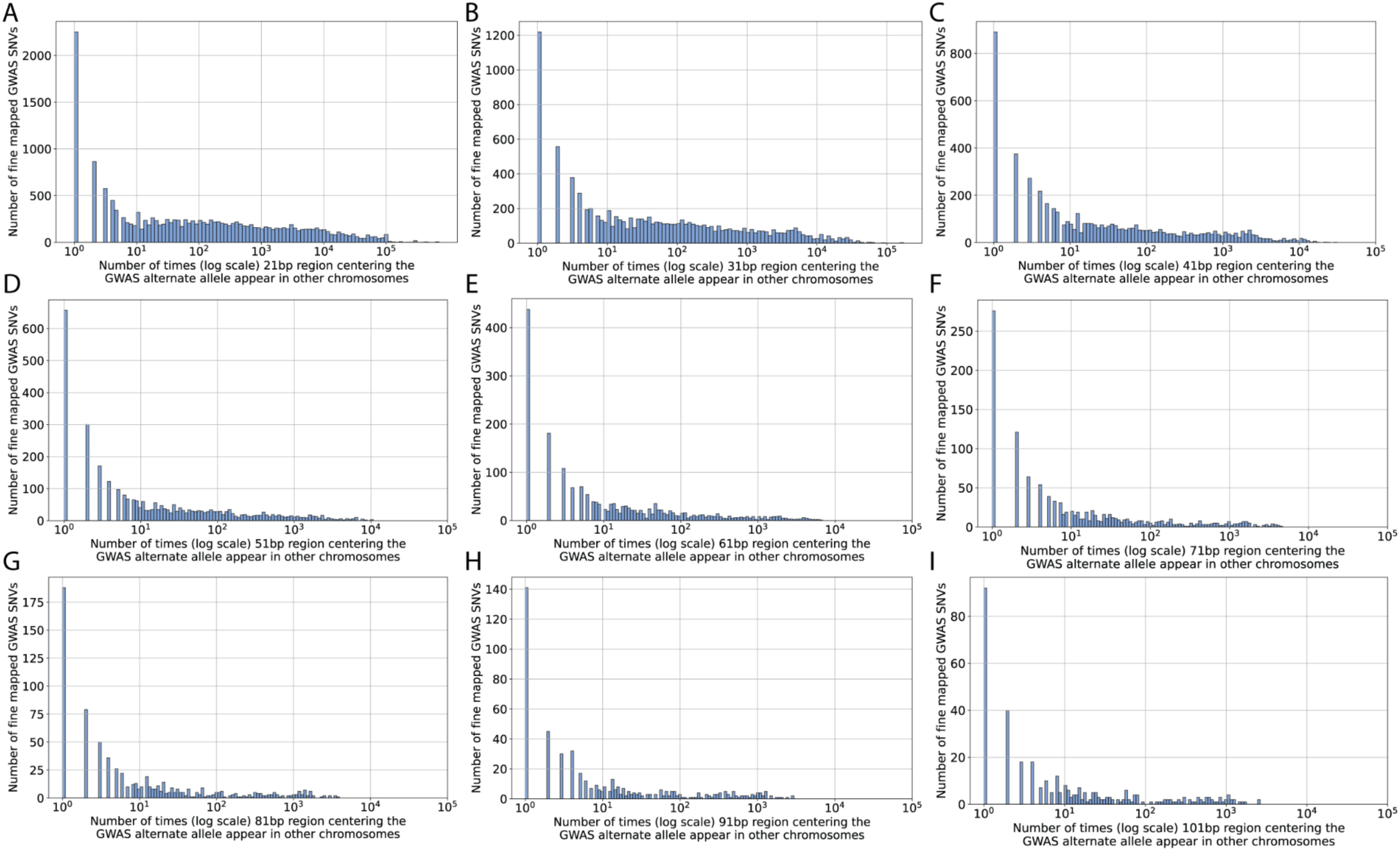
Genome doppelgängers occur on other chromosomes, often many times. Number of fine mapped GWAS SNVs (*y*-axes) with the corresponding number of SNV doppelgängers (*x*-axes) on other chromosomes in the genome for (**A**) 21 bp, (**B**) 31 bp, (**C**) 41 bp, (**D**) 51 bp, (**E**) 61 bp, (**F**) 71 bp, (**G**) 81 bp, (**H**) 91 bp, and (**I**) 101 bp regions.

**Supplementary Figure 6:**
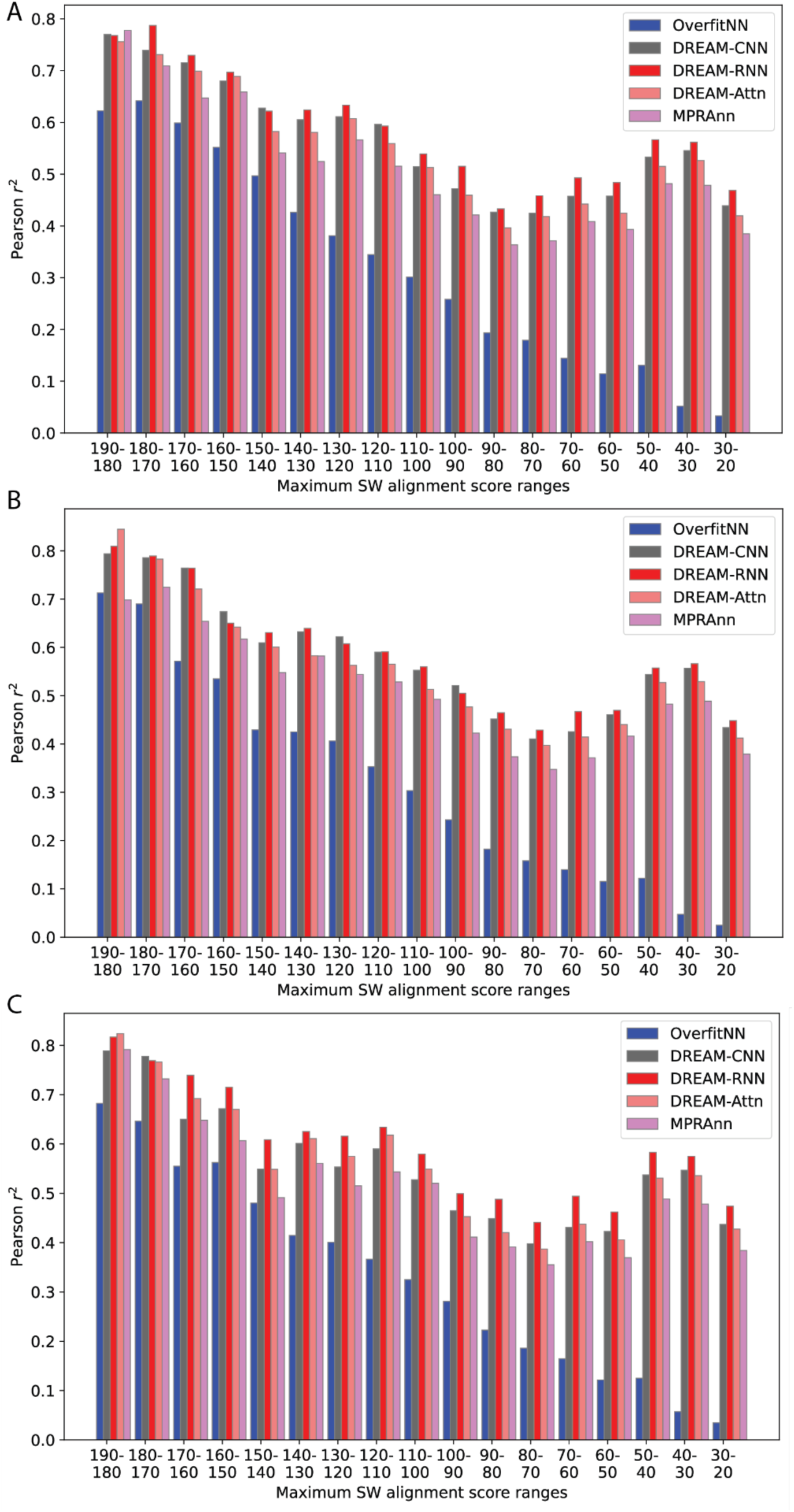
Model performance on test data depends on similarity to training data. Performance comparison (Pearson 𝑟^2^; *y*-axes) of different models (OverfitNN, DREAM-CNN, DREAM-RNN, DREAM-Attn, and MPRAnn; colors) across varying levels of homology (SW alignment score, *x*-axes) in chromosomal fold (**A**) 1, (**B**) 2, and (**C**) 3 (**Methods**).

**Supplementary Figure 7:**
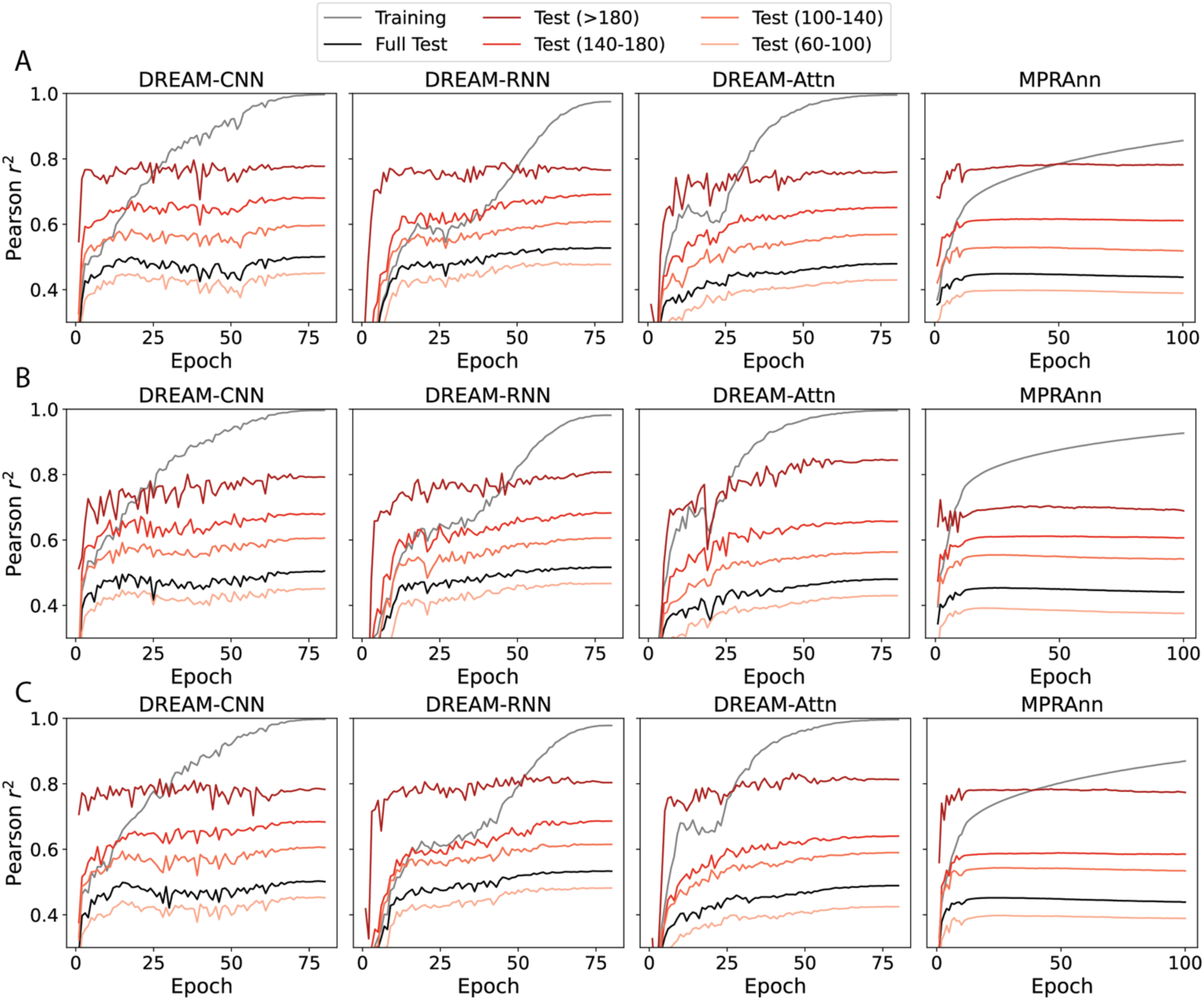
Neural networks trained on three different chromosomal splits show the same trend of varying levels of performance on different degrees of homology. Performance of different models (columns) in Pearson 𝑟^2^(*y*-axes) during model training (*x*-axes) for different sequence sets (colours) for chromosomal spits (A) 1, (B) 2, and (C) 3.

**Supplementary Figure 8:**
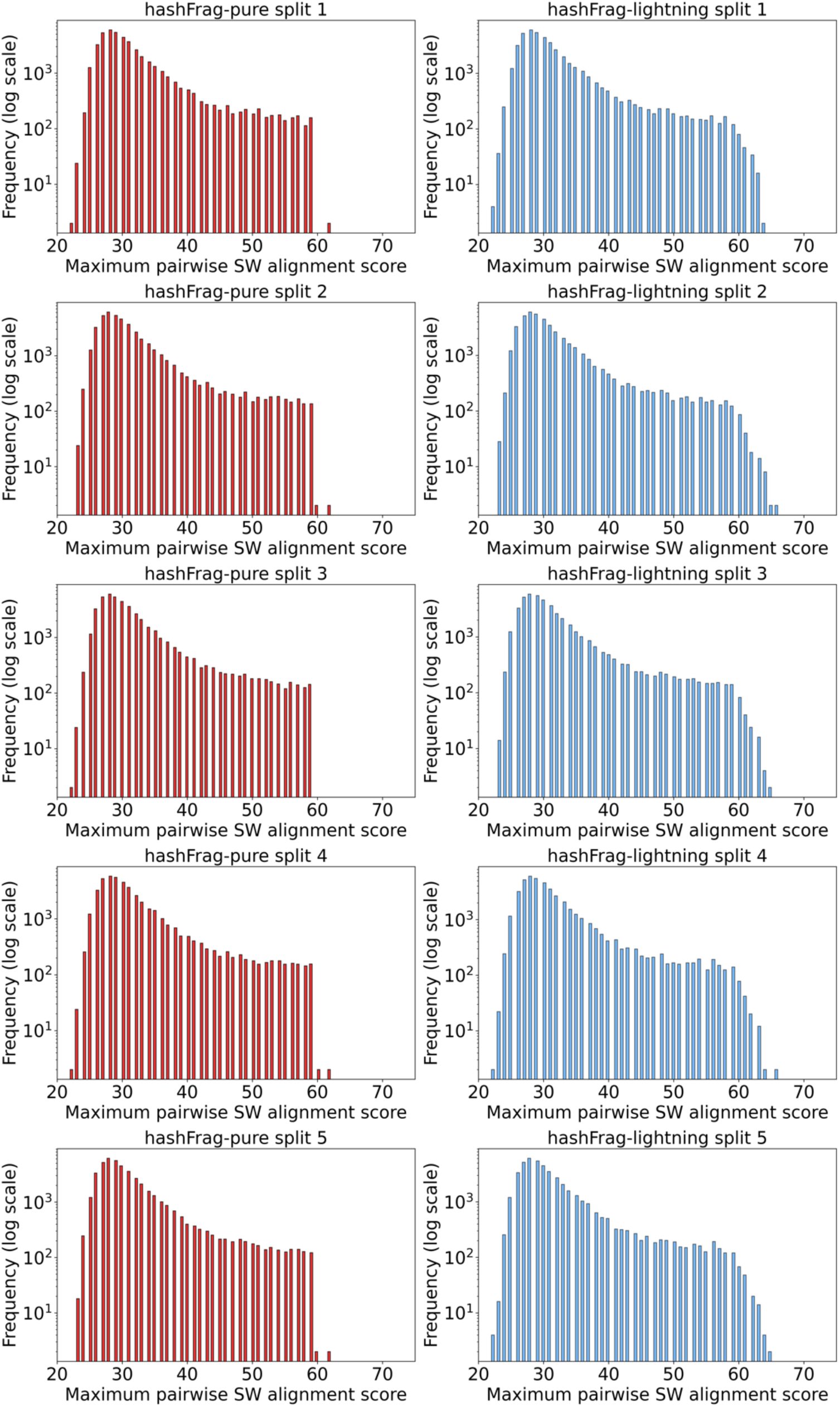
hashFrag-lightning is nearly as effective as hashFrag-pure in detecting homologs. Histogram showing the number of test sequences (*y*-axes) with corresponding maximum pairwise SW local alignment scores with the training sequences (*x*-axis) for different hashFrag-pure (left) and hashFrag-lightning (right) splits (at alignment score threshold of 60). The complete MPRA dataset was used to generate the train-test splits at 80%-20% ratio.

**Supplementary Figure 9:**
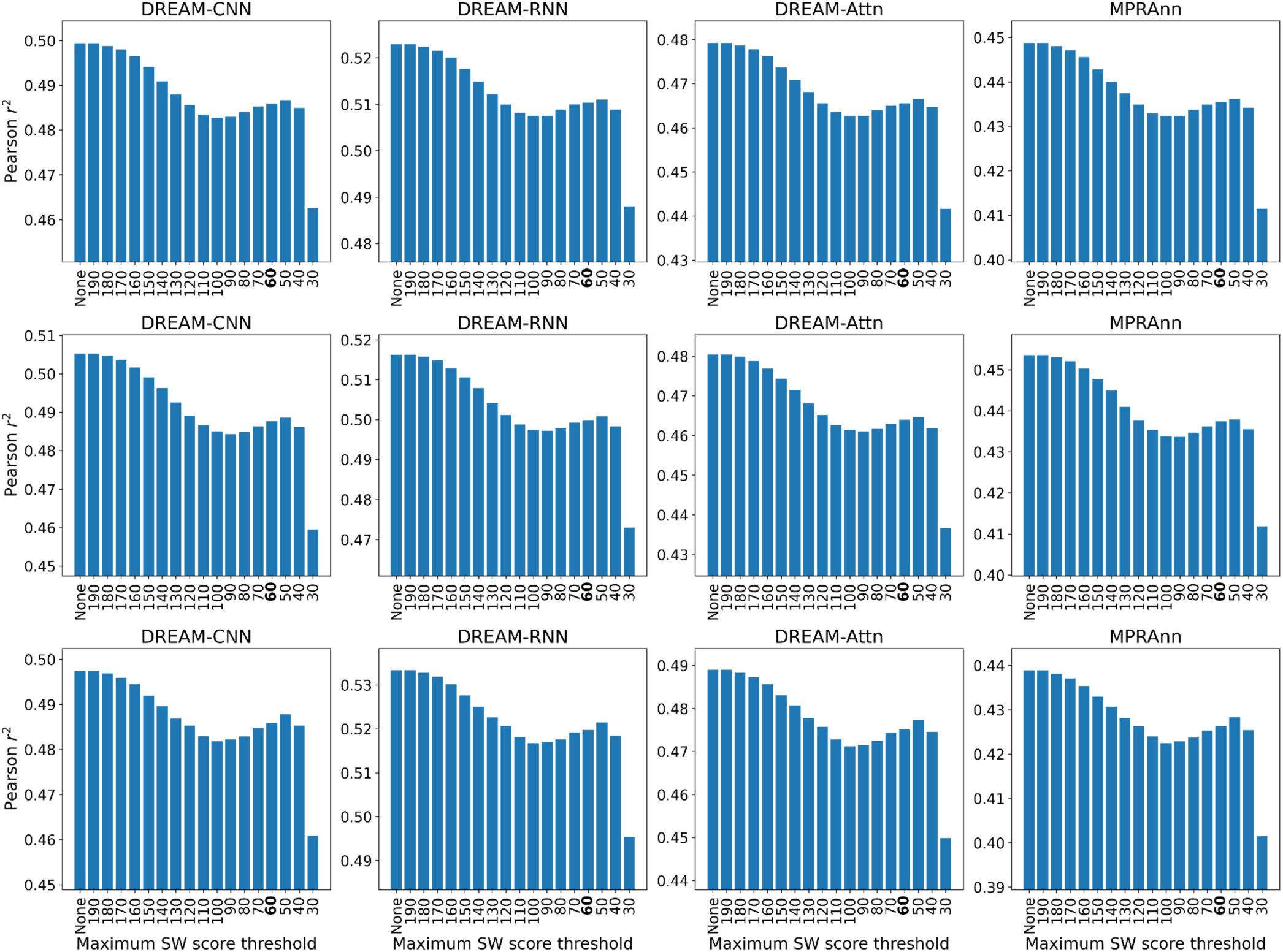
hashFrag removes overestimation of model performance. Model performance (Pearson 𝑟^2^; *y*-axes) across different models (columns) for different chromosomal splits (rows) following the removal of similar sequences using hashFrag-pure at different maximum SW score thresholds (*x*-axes). Models were trained and evaluated on the Agarwal et al. K562 MPRA dataset (39).

**Supplementary Figure 10:**
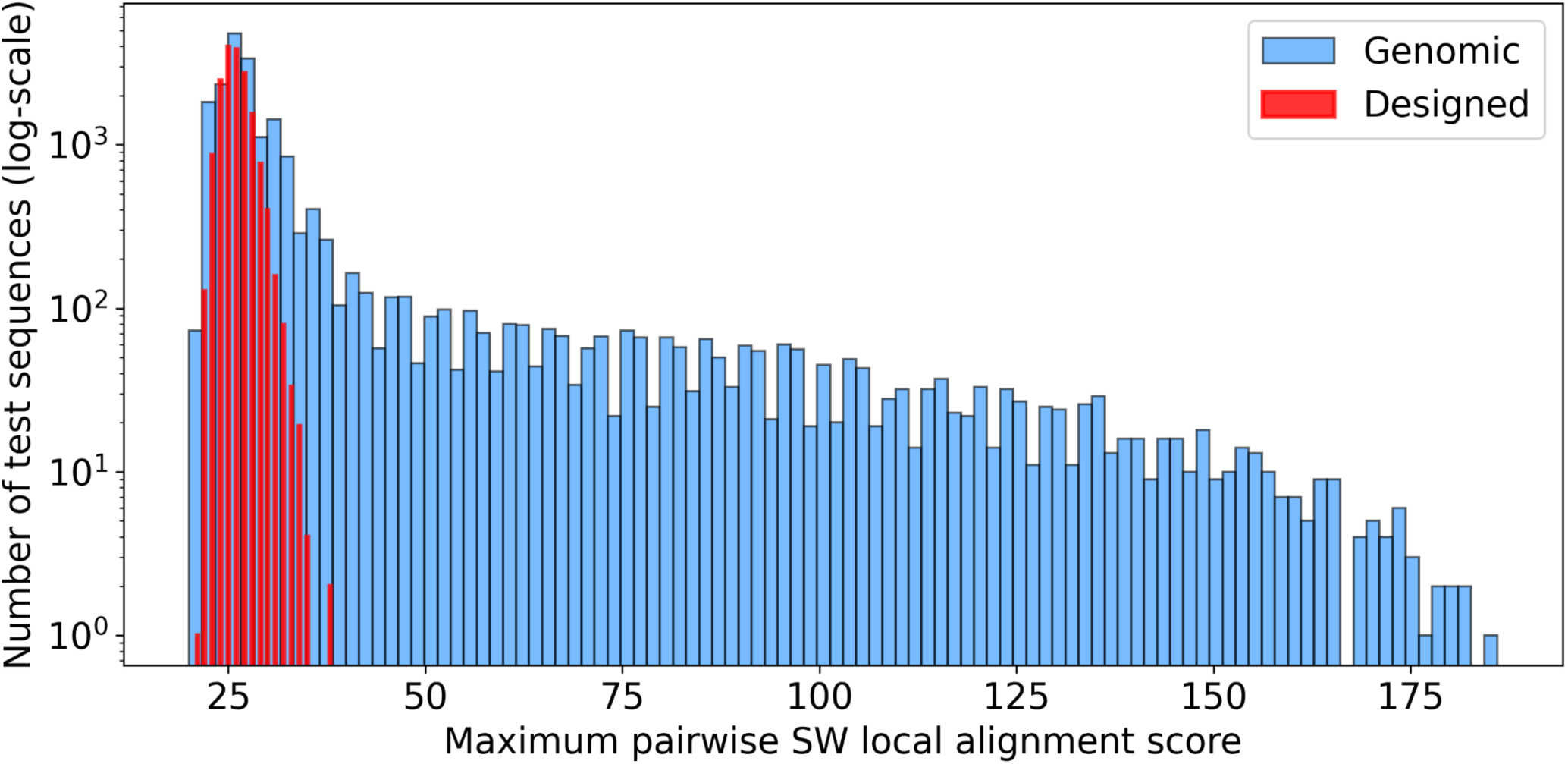
Designed sequences from Gosai et al. (13) are substantially different to genomic sequences (39). Histogram showing the number of test sequences (*y*-axis) with corresponding maximum pairwise SW local alignment scores with the training sequences (*x*-axis). For genomic sequences (blue), training and test sets were randomly sampled from distinct chromosome sets. For designed sequences (red), training sequences included the genome-derived MPRA sequence library and the test sequences were the 17,000 designed sequences from Gosai et al. (13).

**Supplementary Figure 11:**
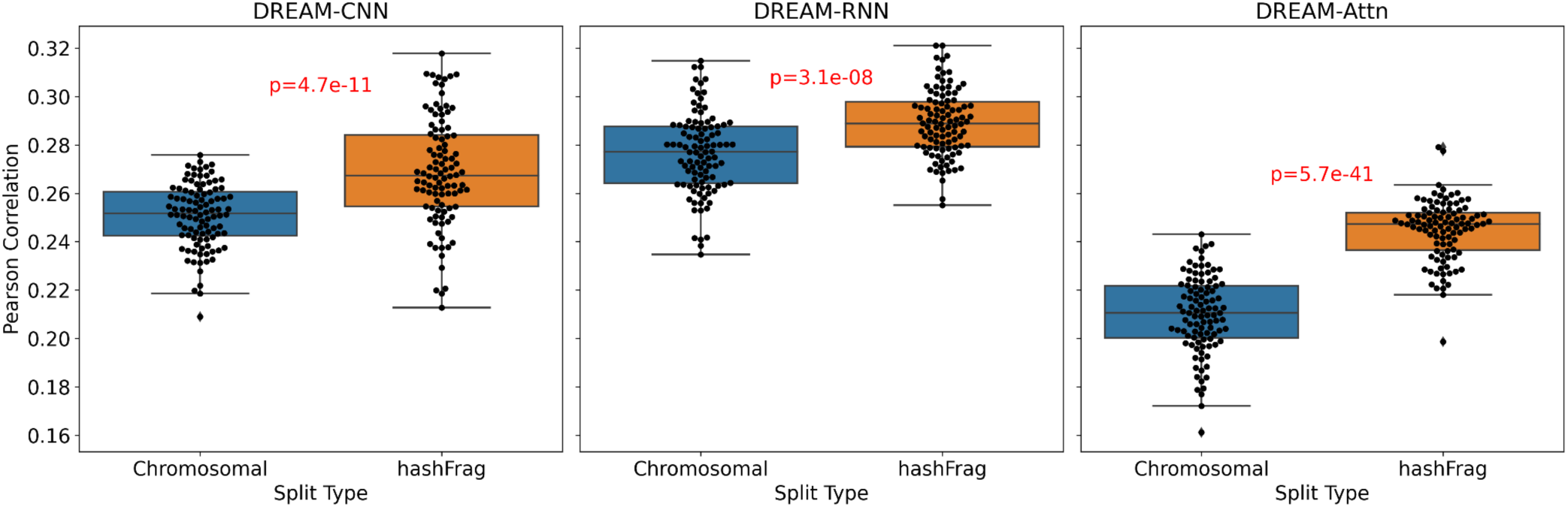
hashFrag-split trained models outperform chromosomal split trained models. Performances (*y*-axes) across 100 replicates (points) of different models (panels) on the designed sequences from Gosai et al. (13) when trained on different chromosomal and hashFrag splits (*x*-axes). Statistical significance between hashFrag and chromosomally trained models was calculated using the Two-Sample t-test.

**Supplementary Figure 12:**
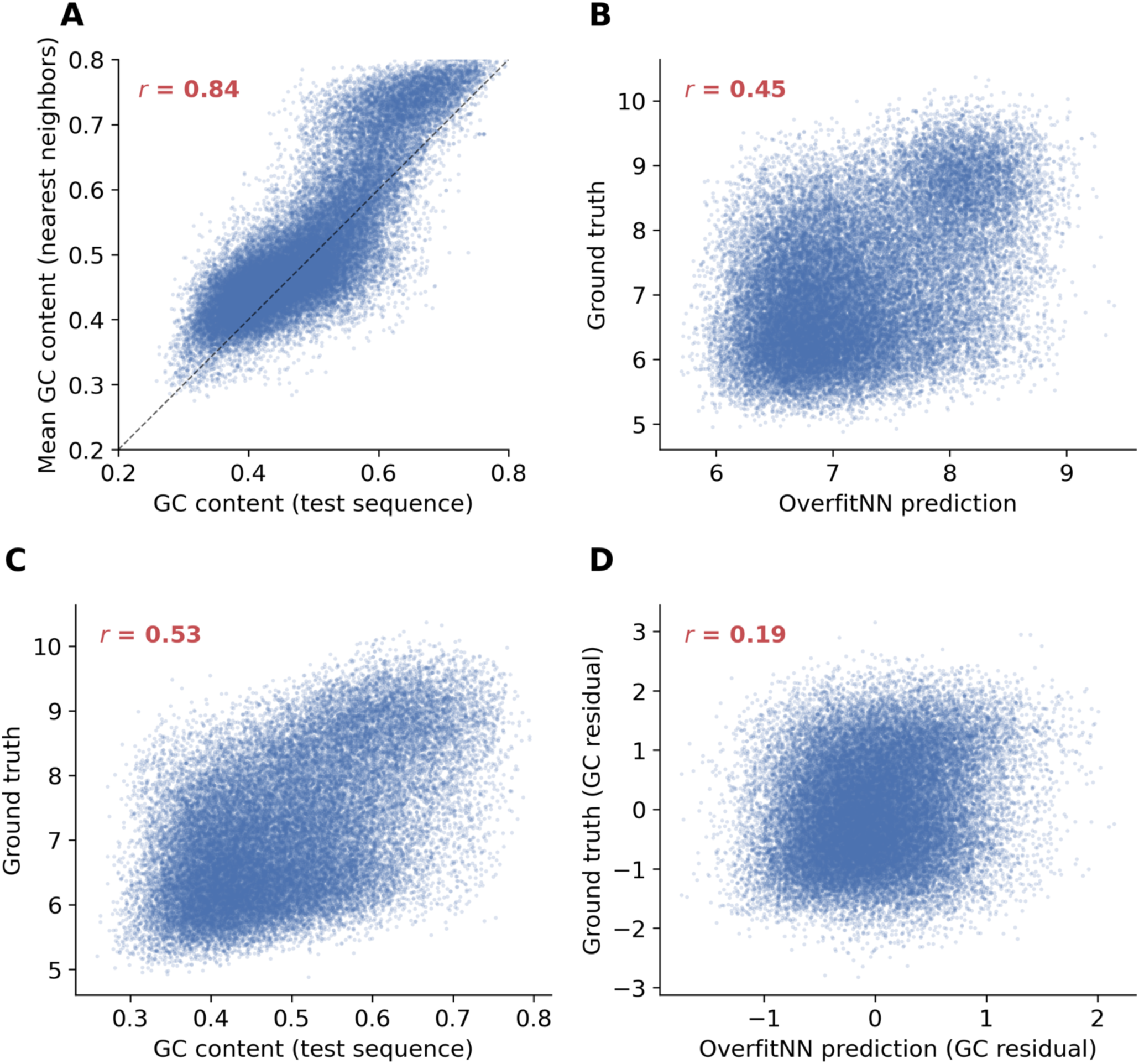
OverfitNN’s performance at low similarity is driven by GC content matching. Analysis restricted to test sequences with maximum SW alignment scores in the 0–100 range (n = 39,880). (**A**) GC content of each test sequence (*x*-axis) versus the mean GC content of its nearest training neighbors (*y*-axis). Neighbors are closely matched to test sequences by GC content. (**B**) OverfitNN predictions (*x*-axis) vs ground truth accessibility read counts (*y*-axis) for each sequence (points). (**C**) GC content (*x*-axis) vs ground truth accessibility read counts (*y*-axis) for each sequence (points) (**D**) OverfitNN predictions (*x*-axis) vs the residual ground truth after regressing out the signal explained by GC content (*y*-axis).

**Supplementary Figure 13:**
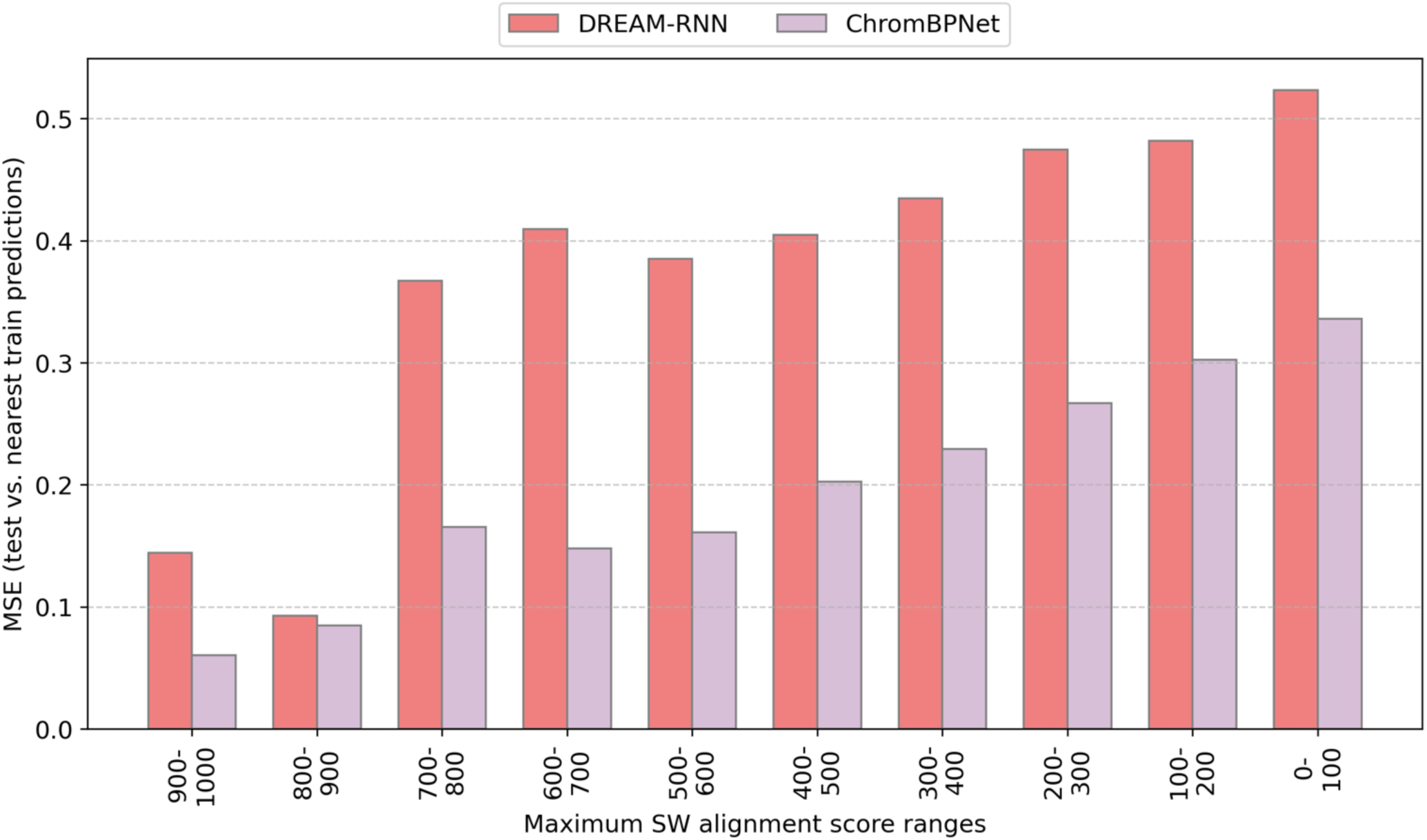
ChromBPNet (33) exhibits higher memorization of similar training sequences compared to DREAM-RNN (66). For each test sequence, MSE (*y*-axis) was computed between the model’s predicted output on the test sequence and its predicted output on the nearest training sequence (highest SW alignment score to the test sequence). Test sequences are binned by maximum SW alignment score to any training sequence (*x*-axis). Lower MSE at high similarity is consistent with memorization. ChromBPNet (color) shows lower MSE than DREAM-RNN (color) across all bins.

**Supplementary Figure 14:**
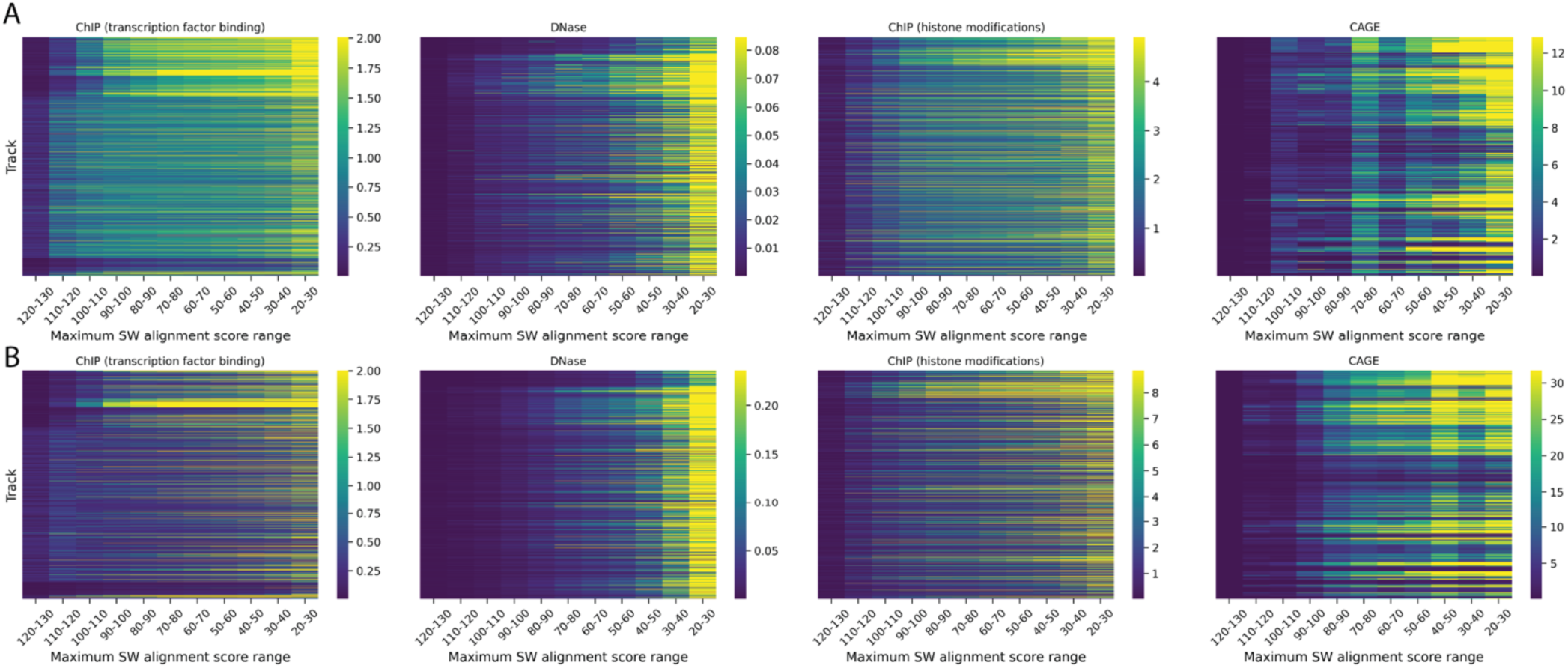
MSE reveals memorization-driven numerical accuracy at high similarity. Heatmaps showing MSE for (**A**) model performance and (**B**) model memorization for individual transcription factor binding ChIP, DNase, histone modifications ChIP, and CAGE tracks (*y*-axes) across bins stratified by maximum SW alignment score (*x*-axes).

**Supplementary Figure 15:**
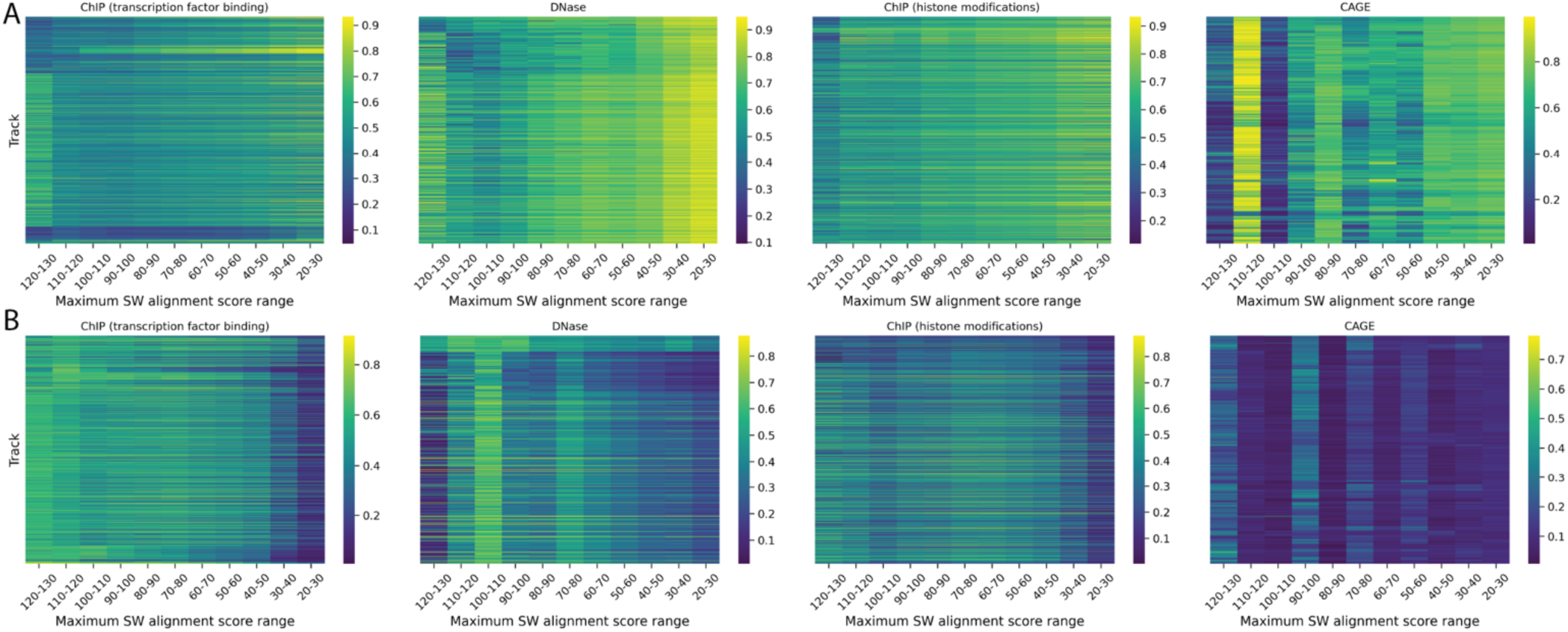
Pearson correlation reveals distinct patterns for model performance and memorization across homology levels. Heatmaps showing Pearson *r* for Enformer (**A**) performance and (**B**) memorization for individual tracks (*y*-axes) for transcription factor binding ChIP, DNase, histone modifications ChIP, and CAGE (subpanels) across bins stratified by maximum SW alignment score (*x*-axes). Tracks are ordered identically by track index in both panels, so each row corresponds to the same track in (**A**) and (**B**) within a given assay subpanel.

**Supplementary Figure 16:**
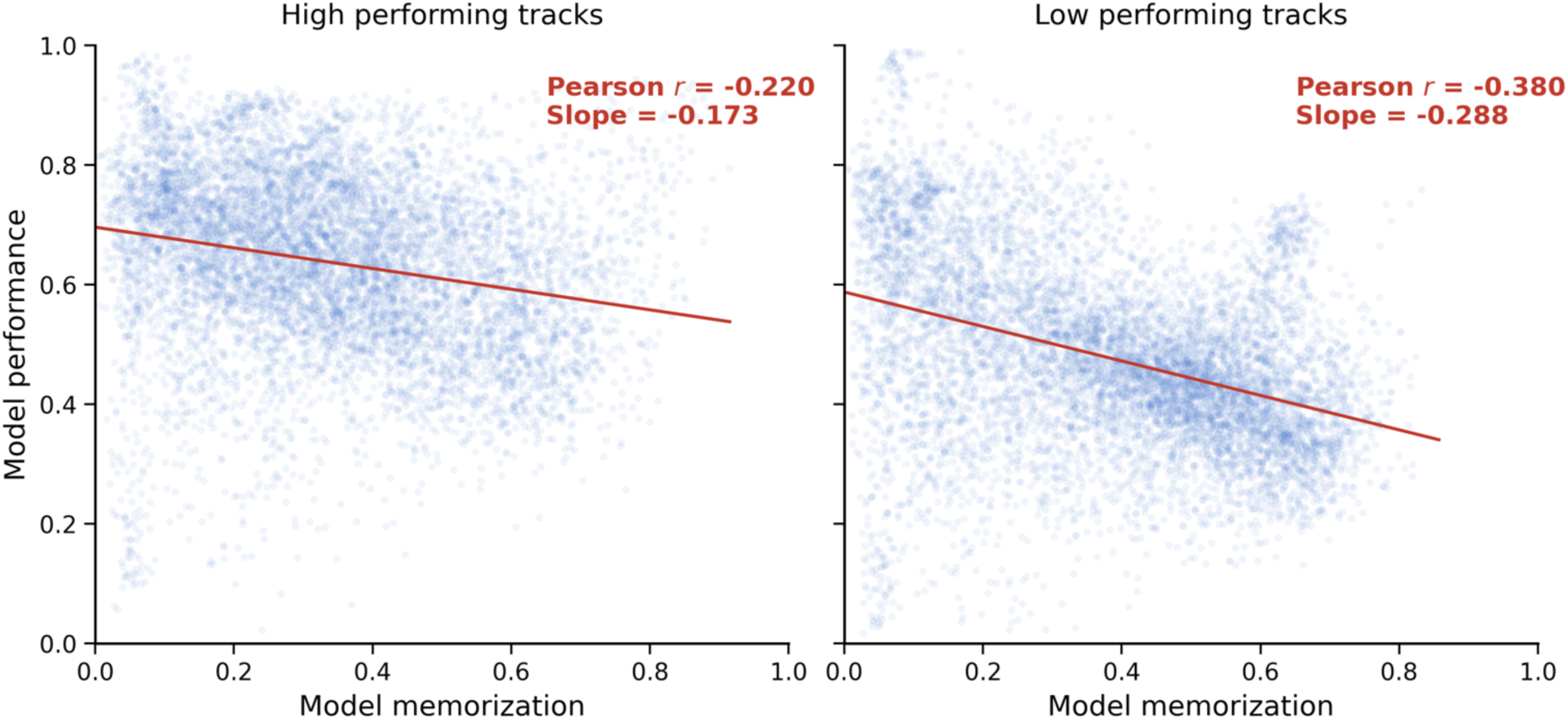
The inverse correlation between memorization and performance is stronger for poorly predicted tracks. Scatter plots showing the relationship between bin-level model performance (Pearson correlation; *x*-axes) and model memorization (Pearson correlation; *y*-axes) across tracks and homology score groups after stratifying Enformer outputs by overall track predictability. Tracks were classified as high performing if their median test-set Pearson correlation was ≥0.5 (**left**) and low performing if their median test-set Pearson correlation was < 0.5 (**right**).

**Supplementary Figure 17:**
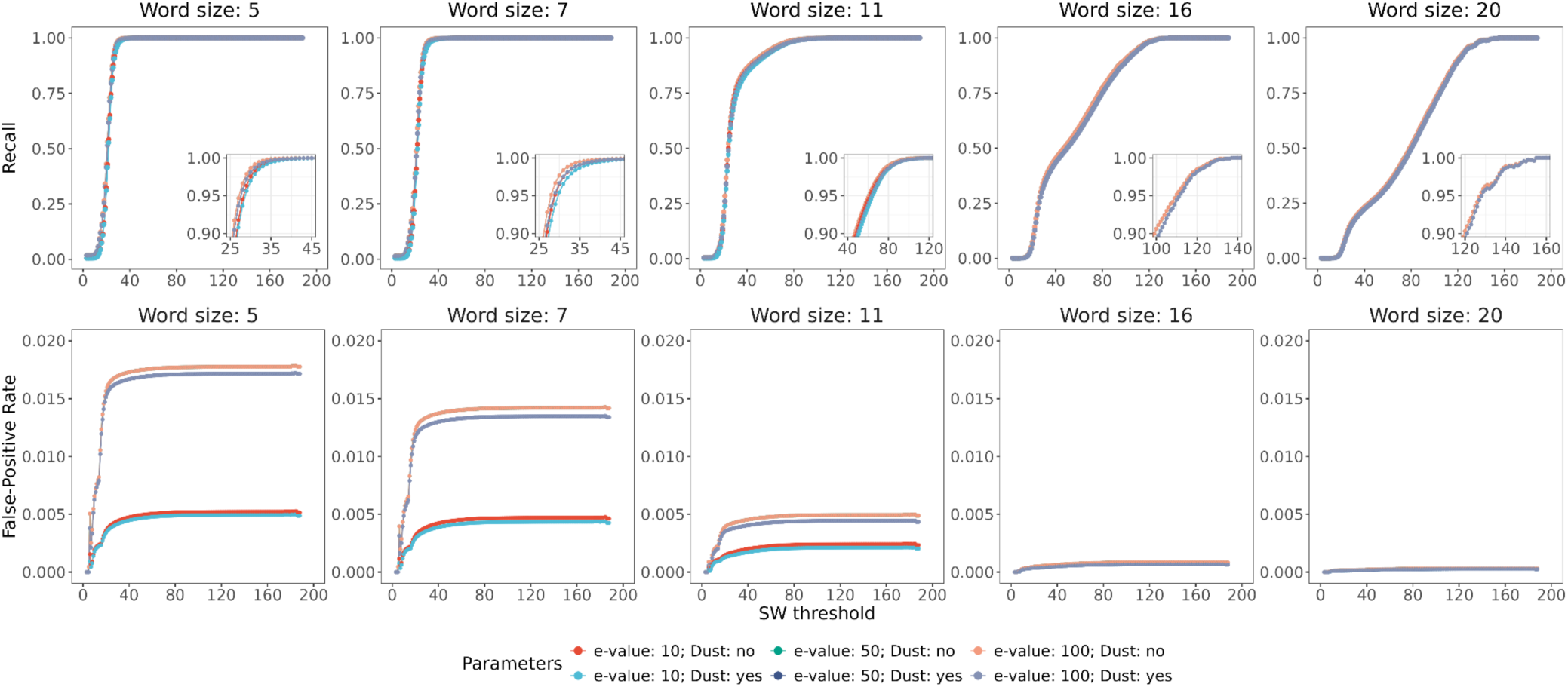
Tuning the BLASTn parameters with respect to recall. 10,000 sequences were randomly sampled from the human K562 MPRA dataset (39) and subjected to the BLASTn algorithm with various parameter combinations (n = 10 replicates) to identify homology within this sequence subset. The following parameters were investigated in terms of recall and false-positive rate (FPR): word_size (5, 7, 11, 16 and 20; columns), max_target_seqs (500, 1000, 5000, and 10000), evalue (10, 50, and 100), and dust (yes, no). Each SW threshold represents the average recall and FPR values across replicates and max_target_seqs parameters. Based on the BLASTn results for each combination of parameters (e-value and dust), recall (*y*-axes top) and FPR (*y*-axes bottom) were calculated for all integer thresholds of SW alignment scores from 0 to 200 (*x*-axes). An inset plot zooming in to the recall results are provided. Notably, no changes in recall or FPR were observed upon marginalization of the max_target_seqs parameter; however this parameter was found to impact recall and FPR for larger datasets.

**Supplementary Figure 18:**
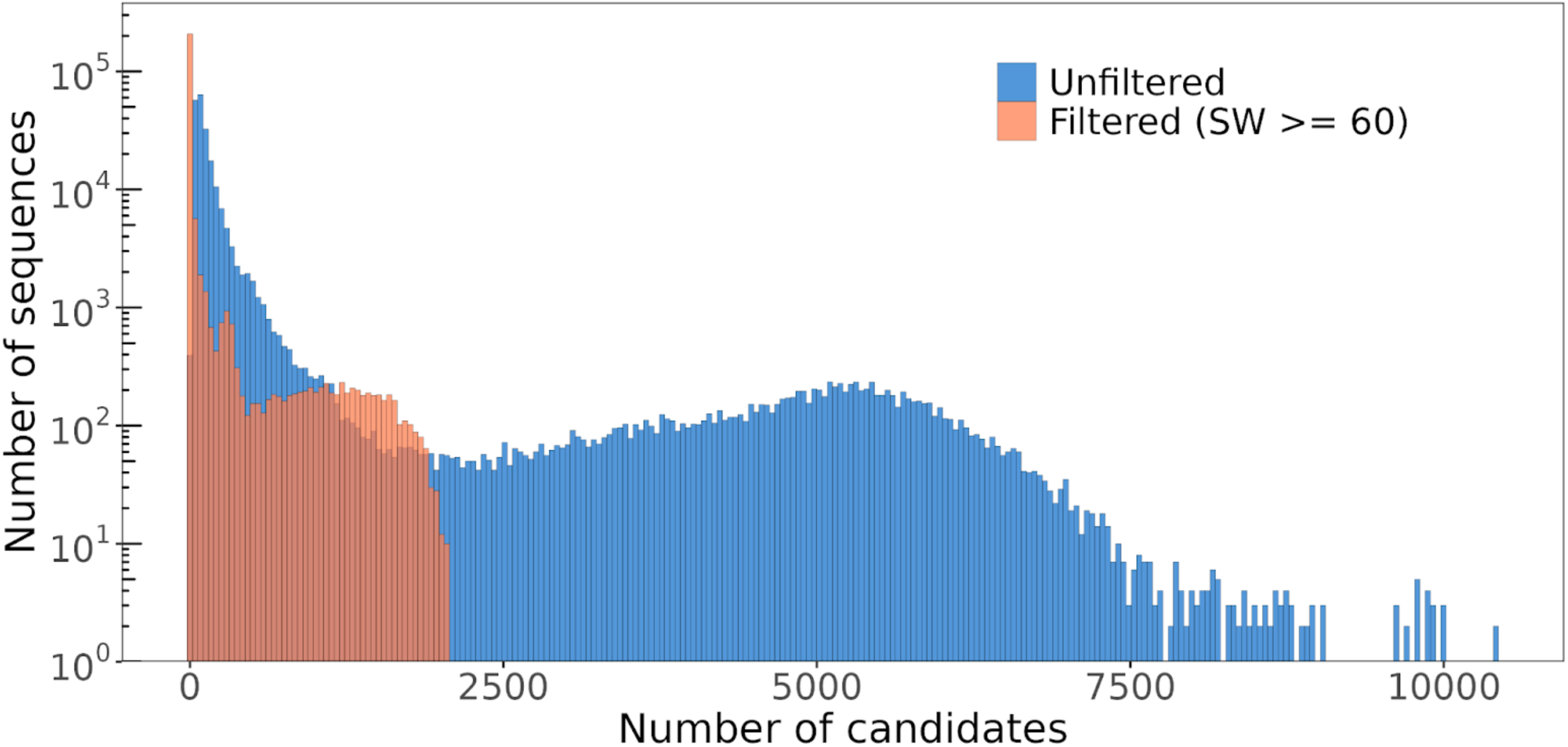
Distribution of the number of candidates identified by hashFrag before and after filtering false-positives by SW score. The BLASTn algorithm was applied to the human K562 MPRA dataset to identify candidates exhibiting similarity (parameters were as follows: word_size=7; match=1, mismatch=-1, gapopen=2, gapextend=1, e_value=100; dust=no; max_target_seqs=226,253). The distribution for the number of candidates initially identified for each sequence is depicted in blue (“Unfiltered”). Based on a SW threshold of 60, we filtered those candidates (shown in orange; “Filtered (SW >= 60)”). In total, approximately 91.1% of identified candidates were filtered based on the defined SW score threshold of 60.

## References

1. Sokolova K, Chen KM, Hao Y, Zhou J, Troyanskaya OG. Deep Learning Sequence Models for Transcriptional Regulation. Annu Rev Genomics Hum Genet. 2024 Aug 27;25(Volume 25, 2024):105–22. doi:10.1146/annurev-genom-021623-024727

2. Sasse A, Chikina M, Mostafavi S. Unlocking gene regulation with sequence-to-function models. Nat Methods. 2024 Aug;21(8):1374–7. doi:10.1038/s41592-024-02331-5

3. Barbadilla-Martínez L, Klaassen N, van Steensel B, de Ridder J. Predicting gene expression from DNA sequence using deep learning models. Nat Rev Genet. 2025 Oct;26(10):666–80. doi:10.1038/s41576-025-00841-2

4. Drusinsky S, Whalen S, Pollard KS. Deep-learning prediction of gene expression from personal genomes. Genome Biol. 2026 Jan 6;27(1):19. doi:10.1186/s13059-025-03926-7

5. Kim S, Wysocka J. Deciphering the multi-scale, quantitative cis-regulatory code. Mol Cell. 2023 Feb 2;83(3):373–92. doi:10.1016/j.molcel.2022.12.032 PubMed PMID: 36693380.

6. Avsec Ž, Weilert M, Shrikumar A, Krueger S, Alexandari A, Dalal K, et al. Base-resolution models of transcription-factor binding reveal soft motif syntax. Nat Genet. 2021 Mar;53(3):3. doi:10.1038/s41588-021-00782-6

7. Tan J, Fu X, Ling X, Mo S, Bai J, Rabadán R, et al. Decoding the gene regulatory landscape through multimodal learning of protein-DNA interactions. bioRxiv; 2026. doi:10.1101/2025.08.17.670761

8. Nair S, Hajiramezanali E, Tseng A, Diamant N, Hingerl J, Lal A, et al. Nona: A unifying multimodal masking framework for functional genomics. bioRxiv; 2025. doi:10.1101/2025.11.06.687036

9. Srivastava D, Mahony S. Sequence and chromatin determinants of transcription factor binding and the establishment of cell type-specific binding patterns. Biochim Biophys Acta BBA - Gene Regul Mech. 2020 Jun 1;Transcriptional Profiles and Regulatory Gene Networks1863(6):194443. doi:10.1016/j.bbagrm.2019.194443

10. Hingerl JC, Martens LD, Karollus A, Manz T, Buenrostro JD, Theis FJ, et al. scooby: modeling multimodal genomic profiles from DNA sequence at single-cell resolution. Nat Methods. 2025 Nov;22(11):2275–85. doi:10.1038/s41592-025-02854-5

11. Dudnyk K, Cai D, Shi C, Xu J, Zhou J. Sequence basis of transcription initiation in the human genome. Science. 2024 Apr 26;384(6694):eadj0116. doi:10.1126/science.adj0116

12. Lal A, Karollus A, Gunsalus L, Garfield D, Nair S, Tseng AM, et al. Decoding sequence determinants of gene expression in diverse cellular and disease states. bioRxiv; 2025 doi:10.1101/2024.10.09.617507

13. Gosai SJ, Castro RI, Fuentes N, Butts JC, Mouri K, Alasoadura M, et al. Machine-guided design of cell-type-targeting cis-regulatory elements. Nature. 2024 Oct;634(8036):1211–20. doi:10.1038/s41586-024-08070-z

14. Taskiran II, Spanier KI, Dickmänken H, Kempynck N, Pančíková A, Ekşi EC, et al. Cell-type-directed design of synthetic enhancers. Nature. 2024 Feb;626(7997):212–20. doi:10.1038/s41586-023-06936-2

15. de Almeida BP, Schaub C, Pagani M, Secchia S, Furlong EEM, Stark A. Targeted design of synthetic enhancers for selected tissues in the Drosophila embryo. Nature. 2024 Feb;626(7997):207–11. doi:10.1038/s41586-023-06905-9

16. DaSilva LF, Senan S, Kribelbauer-Swietek JF, Patel ZM, Louis LK, Reddy AJ, et al. Designing synthetic regulatory elements using the generative AI framework DNA-Diffusion. Nat Genet. 2026 Jan;58(1):180–94. doi:10.1038/s41588-025-02441-6

17. Castillo-Hair SM, Yin CH, VandenBosch L, Cherry TJ, Meuleman W, Seelig G. Programming human cell type-specific gene expression via an atlas of AI-designed enhancers. bioRxiv; 2025. doi:10.1101/2025.09.30.679565

18. Avsec Ž, Latysheva N, Cheng J, Novati G, Taylor KR, Ward T, et al. Advancing regulatory variant effect prediction with AlphaGenome. Nature. 2026 Jan;649(8099):1206–18. doi:10.1038/s41586-025-10014-0

19. Kathail P, Bajwa A, Ioannidis NM. Leveraging genomic deep learning models for the prediction of non-coding variant effects. arXiv; 2025. doi:10.48550/arXiv.2411.11158

20. Butts JC, Rong S, Gosai SJ, Castro RI, Noon M, Adeniran K, et al. Identifying non-coding variant effects at scale via machine learning models of cis-regulatory reporter assays. bioRxiv; 2025. doi:10.1101/2025.04.16.648420

21. Vaz E, Wang L, Galvin J, Keener R, Battle A. Fine-tuning sequence to function deep learning models on large-scale proteomic data improves the accuracy of variant effect prediction. bioRxiv; 2025. doi:10.1101/2025.09.26.678908

22. Jaganathan K, Panagiotopoulou SK, McRae JF, Darbandi SF, Knowles D, Li YI, et al. Predicting Splicing from Primary Sequence with Deep Learning. Cell. 2019 Jan 24;176(3):535–548.e24. doi:10.1016/j.cell.2018.12.015 PubMed PMID: 30661751.

23. Srivastava D, Korsakova A, Wang Q, Ruiz L, Yuan H, Kelley DR. Borzoi-informed fine mapping improves causal variant prioritization in complex trait GWAS. bioRxiv; 2025. doi:10.1101/2025.07.09.663936

24. Zhou J, Park CY, Theesfeld CL, Wong AK, Yuan Y, Scheckel C, et al. Whole-genome deep-learning analysis identifies contribution of noncoding mutations to autism risk. Nat Genet. 2019 Jun;51(6):973–80. doi:10.1038/s41588-019-0420-0

25. Chen KM, Wong AK, Troyanskaya OG, Zhou J. A sequence-based global map of regulatory activity for deciphering human genetics. Nat Genet. 2022 Jul;54(7):940–9. doi:10.1038/s41588-022-01102-2

26. Naqvi S, Kim S, Tabatabaee S, Pampari A, Kundaje A, Pritchard JK, et al. Transfer learning reveals sequence determinants of the quantitative response to transcription factor dosage. Cell Genomics. 2025 Mar 12;5(3). doi:10.1016/j.xgen.2025.100780 PubMed PMID: 40020686.

27. de Boer CG, Taipale J. Hold out the genome: a roadmap to solving the cis-regulatory code. Nature. 2024 Jan;625(7993):41–50. doi:10.1038/s41586-023-06661-w

28. Friedman RZ, Ramu A, Lichtarge S, Wu Y, Tripp L, Lyon D, et al. Active learning of enhancers and silencers in the developing neural retina. Cell Syst. 2025 Jan;16(1):101163. doi:10.1016/j.cels.2024.12.004

29. Luthra I, Jensen C, Chen XE, Salaudeen AL, Rafi AM, de Boer CG. Regulatory activity is the default DNA state in eukaryotes. Nat Struct Mol Biol. 2024 Mar;31(3):559–67. doi:10.1038/s41594-024-01235-4

30. Linder J, Srivastava D, Yuan H, Agarwal V, Kelley DR. Predicting RNA-seq coverage from DNA sequence as a unifying model of gene regulation. Nat Genet. 2025 Apr;57(4):949–61. doi:10.1038/s41588-024-02053-6

31. Avsec Ž, Agarwal V, Visentin D, Ledsam JR, Grabska-Barwinska A, Taylor KR, et al. Effective gene expression prediction from sequence by integrating long-range interactions. Nat Methods. 2021 Oct;18(10):10. doi:10.1038/s41592-021-01252-x

32. Cazares TA, Rizvi FW, Iyer B, Chen X, Kotliar M, Bejjani AT, et al. maxATAC: Genome-scale transcription-factor binding prediction from ATAC-seq with deep neural networks. PLOS Comput Biol. 2023 Jan 31;19(1):e1010863. doi:10.1371/journal.pcbi.1010863

33. Pampari A, Shcherbina A, Kvon EZ, Kosicki M, Nair S, Kundu S, et al. ChromBPNet: bias factorized, base-resolution deep learning models of chromatin accessibility reveal cis-regulatory sequence syntax, transcription factor footprints and regulatory variants. bioRxiv; 2025. doi:10.1101/2024.12.25.630221

34. Jung AJ, Zhu H, Gao AJ, Li R, Slobodyanyuk M, Chu V, et al. FlashRNA: An Efficient Model for Regulatory Genomics. bioRxiv; 2025. doi:10.1101/2025.10.14.682350

35. Celaj A, Gao AJ, Lau TTY, Holgersen EM, Lo A, Lodaya V, et al. An RNA foundation model enables discovery of disease mechanisms and candidate therapeutics. bioRxiv; 2023. doi:10.1101/2023.09.20.558508

36. Hingerl JC, Karollus A, Gagneur J. Flashzoi: an enhanced Borzoi for accelerated genomic analysis. Bioinformatics. 2025 Sep 1;41(9):btaf467. doi:10.1093/bioinformatics/btaf467

37. de Almeida BP, Reiter F, Pagani M, Stark A. DeepSTARR predicts enhancer activity from DNA sequence and enables the de novo design of synthetic enhancers. Nat Genet. 2022 May;54(5):5. doi:10.1038/s41588-022-01048-5

38. Sahu B, Hartonen T, Pihlajamaa P, Wei B, Dave K, Zhu F, et al. Sequence determinants of human gene regulatory elements. Nat Genet. 2022 Mar;54(3):3. doi:10.1038/s41588-021-01009-4

39. Agarwal V, Inoue F, Schubach M, Penzar D, Martin BK, Dash PM, et al. Massively parallel characterization of transcriptional regulatory elements. Nature. 2025 Mar;639(8054):411–20. doi:10.1038/s41586-024-08430-9

40. Penzar D, Nogina D, Noskova E, Zinkevich A, Meshcheryakov G, Lando A, et al. LegNet: a best-in-class deep learning model for short DNA regulatory regions. Bioinformatics. 2023 Aug 1;39(8):btad457. doi:10.1093/bioinformatics/btad457

41. Koning APJ de, Gu W, Castoe TA, Batzer MA, Pollock DD. Repetitive Elements May Comprise Over Two-Thirds of the Human Genome. PLOS Genet. 2011 Dec 1;7(12):e1002384. doi:10.1371/journal.pgen.1002384

42. Lander ES, Linton LM, Birren B, Nusbaum C, Zody MC, Baldwin J, et al. Initial sequencing and analysis of the human genome. Nature. 2001 Feb;409(6822):860–921. doi:10.1038/35057062

43. Whalen S, Schreiber J, Noble WS, Pollard KS. Navigating the pitfalls of applying machine learning in genomics. Nat Rev Genet. 2022 Mar;23(3):169–81. doi:10.1038/s41576-021-00434-9

44. Bernett J, Blumenthal DB, Grimm DG, Haselbeck F, Joeres R, Kalinina OV, et al. Guiding questions to avoid data leakage in biological machine learning applications. Nat Methods. 2024 Aug;21(8):1444–53. doi:10.1038/s41592-024-02362-y

45. Joeres R, Blumenthal DB, Kalinina OV. Data splitting to avoid information leakage with DataSAIL. Nat Commun. 2025 Apr 8;16(1):3337. doi:10.1038/s41467-025-58606-8

46. Kapoor S, Narayanan A. Leakage and the reproducibility crisis in machine-learning-based science. Patterns. 2023 Sep 8;4(9):100804. doi:10.1016/j.patter.2023.100804

47. Rafi AM, Nogina D, Penzar D, Lee D, Lee D, Kim N, et al. A community effort to optimize sequence-based deep learning models of gene regulation. Nat Biotechnol. 2025 Aug;43(8):1373–83. doi:10.1038/s41587-024-02414-w

48. Altschul SF, Gish W, Miller W, Myers EW, Lipman DJ. Basic local alignment search tool. J Mol Biol. 1990 Oct 5;215(3):403–10. doi:10.1016/S0022-2836(05)80360-2

49. Camacho C, Coulouris G, Avagyan V, Ma N, Papadopoulos J, Bealer K, et al. BLAST+: architecture and applications. BMC Bioinformatics. 2009 Dec 15;10(1):421. doi:10.1186/1471-2105-10-421

50. Arpit D, Jastrzębski S, Ballas N, Krueger D, Bengio E, Kanwal MS, et al. A Closer Look at Memorization in Deep Networks. arXiv; 2017. doi:10.48550/arXiv.1706.05394

51. Zhang C, Bengio S, Hardt M, Recht B, Vinyals O. Understanding deep learning requires rethinking generalization. arXiv; 2017. doi:10.48550/arXiv.1611.03530

52. Ferrer Florensa A, Almagro Armenteros JJ, Nielsen H, Aarestrup FM, Clausen PTLC. SpanSeq: similarity-based sequence data splitting method for improved development and assessment of deep learning projects. NAR Genomics Bioinforma. 2024 Sep 1;6(3):lqae106. doi:10.1093/nargab/lqae106

53. Cawley GC, Talbot NLC. On Over-fitting in Model Selection and Subsequent Selection Bias in Performance Evaluation. J Mach Learn Res. 2010;11(70):2079–107.

54. Smith TF, Waterman MS. Identification of common molecular subsequences. J Mol Biol. 1981 Mar 25;147(1):195–7. doi:10.1016/0022-2836(81)90087-5

55. Gordân R, Narlikar L, Hartemink AJ. Finding regulatory DNA motifs using alignment-free evolutionary conservation information. Nucleic Acids Res. 2010 Apr 1;38(6):e90. doi:10.1093/nar/gkp1166

56. Mountjoy E, Schmidt EM, Carmona M, Schwartzentruber J, Peat G, Miranda A, et al. An open approach to systematically prioritize causal variants and genes at all published human GWAS trait-associated loci. Nat Genet. 2021 Nov;53(11):1527–33. doi:10.1038/s41588-021-00945-5

57. Sasse A, Ng B, Spiro AE, Tasaki S, Bennett DA, Gaiteri C, et al. Benchmarking of deep neural networks for predicting personal gene expression from DNA sequence highlights shortcomings. Nat Genet. 2023 Dec;55(12):2060–4. doi:10.1038/s41588-023-01524-6

58. Huang C, Shuai RW, Baokar P, Chung R, Rastogi R, Kathail P, et al. Personal transcriptome variation is poorly explained by current genomic deep learning models. Nat Genet. 2023 Dec;55(12):2056–9. doi:10.1038/s41588-023-01574-w

59. Benegas G, Batra SS, Song YS. DNA language models are powerful predictors of genome-wide variant effects. Proc Natl Acad Sci. 2023 Oct 31;120(44):e2311219120. doi:10.1073/pnas.2311219120

60. Mclaughlin SM, Lim DA. Probing genomic language models: Nucleotide Generative Pretrained Transformer and the role of pretraining in learned representations. Brief Bioinform. 2026 Jan 1;27(1):bbag011. doi:10.1093/bib/bbag011

61. Kim T, Shin J, Kim H, Jung Y, Lee J, Lee WC, et al. DNACHUNKER: Learnable Tokenization for DNA Language Models. arXiv; 2026. doi:10.48550/arXiv.2601.03019

62. Nguyen E, Poli M, Durrant MG, Kang B, Katrekar D, Li DB, et al. Sequence modeling and design from molecular to genome scale with Evo. Science. 2024 Nov 15;386(6723):eado9336. doi:10.1126/science.ado9336

63. Chuong EB, Elde NC, Feschotte C. Regulatory activities of transposable elements: from conflicts to benefits. Nat Rev Genet. 2017 Feb;18(2):71–86. doi:10.1038/nrg.2016.139

64. Jacques PÉ, Jeyakani J, Bourque G. The Majority of Primate-Specific Regulatory Sequences Are Derived from Transposable Elements. PLOS Genet. 2013 May 9;9(5):e1003504. doi:10.1371/journal.pgen.1003504

65. Sundaram V, Cheng Y, Ma Z, Li D, Xing X, Edge P, et al. Widespread contribution of transposable elements to the innovation of gene regulatory networks. Genome Res. 2014 Jan 12;24(12):1963–76. doi:10.1101/gr.168872.113 PubMed PMID: 25319995.

66. Rafi AM, Nogina D, Penzar D, Lee D, Lee D, Kim N, et al. A community effort to optimize sequence-based deep learning models of gene regulation. Nat Biotechnol. 2025 Aug;43(8):1373–83. doi:10.1038/s41587-024-02414-w

67. Inoue F, Ahituv N. Decoding enhancers using massively parallel reporter assays. Genomics. 2015 Sep 1;Recent advances in functional assays of transcriptional enhancers106(3):159–64. doi:10.1016/j.ygeno.2015.06.005

68. Klein JC, Agarwal V, Inoue F, Keith A, Martin B, Kircher M, et al. A systematic evaluation of the design and context dependencies of massively parallel reporter assays. Nat Methods. 2020 Nov;17(11):1083–91. doi:10.1038/s41592-020-0965-y

69. Ordoñez R, Zhang W, Ellis G, Zhu Y, Ashe HJ, Ribeiro-dos-Santos AM, et al. Genomic context sensitizes regulatory elements to genetic disruption. Mol Cell. 2024 May 16;84(10):1842–1854.e7. doi:10.1016/j.molcel.2024.04.013 PubMed PMID: 38759624.

70. Kim S, Wysocka J. Deciphering the multi-scale, quantitative cis-regulatory code. Mol Cell. 2023 Feb 2;83(3):373–92. doi:10.1016/j.molcel.2022.12.032 PubMed PMID: 36693380.

71. Hammelman J, Krismer K, Banerjee B, Gifford DK, Sherwood RI. Identification of determinants of differential chromatin accessibility through a massively parallel genome-integrated reporter assay. Genome Res. 2020 Jan 10;30(10):1468–80. doi:10.1101/gr.263228.120 PubMed PMID: 32973041.

72. Agarwal V, Shendure J. Predicting mRNA Abundance Directly from Genomic Sequence Using Deep Convolutional Neural Networks. Cell Rep. 2020 May 19;31(7). doi:10.1016/j.celrep.2020.107663 PubMed PMID: 32433972.

73. de Boer CG, Vaishnav ED, Sadeh R, Abeyta EL, Friedman N, Regev A. Deciphering eukaryotic gene-regulatory logic with 100 million random promoters. Nat Biotechnol. 2020 Jan;38(1):1. doi:10.1038/s41587-019-0315-8

74. Karollus A, Mauermeier T, Gagneur J. Current sequence-based models capture gene expression determinants in promoters but mostly ignore distal enhancers. Genome Biol. 2023 Mar 27;24(1):56. doi:10.1186/s13059-023-02899-9

75. Dunham I, Kundaje A, Aldred SF, Collins PJ, Davis CA, Doyle F, et al. An integrated encyclopedia of DNA elements in the human genome. Nature. 2012 Sep;489(7414):57–74. doi:10.1038/nature11247

76. Andersson R, Gebhard C, Miguel-Escalada I, Hoof I, Bornholdt J, Boyd M, et al. An atlas of active enhancers across human cell types and tissues. Nature. 2014 Mar;507(7493):455–61. doi:10.1038/nature12787

77. Treangen TJ, Salzberg SL. Repetitive DNA and next-generation sequencing: computational challenges and solutions. Nat Rev Genet. 2012 Jan;13(1):36–46. doi:10.1038/nrg3117

78. Alser M, Rotman J, Deshpande D, Taraszka K, Shi H, Baykal PI, et al. Technology dictates algorithms: recent developments in read alignment. Genome Biol. 2021 Aug 26;22(1):249. doi:10.1186/s13059-021-02443-7

79. Morrissey A, Shi J, James DQ, Mahony S. Accurate allocation of multimapped reads enables regulatory element analysis at repeats. Genome Res. 2024 Jun;34(6):937–51. doi:10.1101/gr.278638.123 PubMed PMID: 38986578; PubMed Central PMCID: PMC11293539.

80. Cazottes E, Alfeghaly C, Rognard C, Necsulea A, Loda A, Castel G, et al. Remodeling of XIST regulatory landscape during primate evolution. Sci Adv. 2026 Jan 16;12(3):eadw5839. doi:10.1126/sciadv.adw5839

81. Lu AX, Lu AX, Moses A. Evolution Is All You Need: Phylogenetic Augmentation for Contrastive Learning. arXiv; 2020. doi:10.48550/arXiv.2012.13475

82. Lee NK, Tang Z, Toneyan S, Koo PK. EvoAug: improving generalization and interpretability of genomic deep neural networks with evolution-inspired data augmentations. Genome Biol. 2023 May 5;24(1):105. doi:10.1186/s13059-023-02941-w

83. Duncan AG, Mitchell JA, Moses AM. Improving the performance of supervised deep learning for regulatory genomics using phylogenetic augmentation. Bioinformatics. 2024 Apr 1;40(4):btae190. doi:10.1093/bioinformatics/btae190

84. Qiu C, Daza RM, Welsh IC, Patwardhan RP, Martin BK, Li T, et al. Evolutionary transfer learning enables organism-wide inference of mammalian enhancer landscapes. bioRxiv; 2026 doi:10.64898/2026.04.07.717039

85. Vaishnav ED, de Boer CG, Molinet J, Yassour M, Fan L, Adiconis X, et al. The evolution, evolvability and engineering of gene regulatory DNA. Nature. 2022 Mar;603(7901):7901. doi:10.1038/s41586-022-04506-6

86. Sarkar A, Duran A, Yu Y, Lin DW, Kang Y, Somia N, et al. Designing DNA With Tunable Regulatory Activity Using Discrete Diffusion. bioRxiv; 2026. doi:10.1101/2024.05.23.595630

87. Cock PJA, Antao T, Chang JT, Chapman BA, Cox CJ, Dalke A, et al. Biopython: freely available Python tools for computational molecular biology and bioinformatics. Bioinformatics. 2009 Jun 1;25(11):1422–3. doi:10.1093/bioinformatics/btp163

